# Quantitative trait loci mapping of circulating metabolites in cerebrospinal fluid to uncover biological mechanisms involved in brain-related phenotypes

**DOI:** 10.1101/2023.09.26.559021

**Authors:** Lianne M. Reus, Toni Boltz, Marcelo Francia, Merel Bot, Naren Ramesh, Maria Koromina, Yolande A.L. Pijnenburg, Anouk den Braber, Wiesje M. van der Flier, Pieter Jelle Visser, Sven J. van der Lee, Betty M. Tijms, Charlotte E. Teunissen, Loes Olde Loohuis, Roel A. Ophoff

**Affiliations:** Center for Neurobehavioral Genetics, Semel Institute for Neuroscience and Human Behavior, University of California, Los Angeles, CA, USA; Alzheimer Center Amsterdam, Neurology, Vrije Universiteit Amsterdam, Amsterdam UMC location VUmc, Amsterdam, The Netherlands; Amsterdam Neuroscience, Neurodegeneration, Vrije Universiteit Amsterdam, Amsterdam UMC location VUmc, Amsterdam, The Netherlands; Department of Human Genetics, David Geffen School of Medicine, University of California Los Angeles, Los Angeles, CA, USA; Charles Bronfman Institute for Personalized Medicine, Icahn School of Medicine at Mount Sinai, New York, New York, Department of Psychiatry, Icahn School of Medicine at Mount Sinai, New York, New York, Department of Genetics and Genomic Sciences, Icahn School of Medicine at Mount Sinai, New York, New York; Department of Biological Psychology, Amsterdam Public Health Research Institute, Vrije Universiteit Amsterdam, Amsterdam, The Netherlands; Department of Psychiatry, Maastricht University, Maastricht, The Netherlands; Department of Neurobiology, Care Sciences and Society, Division of Neurogeriatrics, Karolinska Institutet, Stockholm, Sweden; Section Genomics of Neurodegenerative Diseases and Aging, Department of Clinical Genetics, Vrije Universiteit Amsterdam, Amsterdam UMC, Amsterdam, the Netherlands; Neurochemistry Lab and Biobank, Department of Clinical Chemistry, Amsterdam Neuroscience, Amsterdam UMC, Amsterdam, The Netherlands; Department of Psychiatry, Erasmus University Medical Center, Rotterdam, The Netherlands

## Abstract

Genomic studies of molecular traits have provided mechanistic insights into complex disease, though these lag behind for brain-related traits due to the inaccessibility of brain tissue. We leveraged cerebrospinal fluid (CSF) to study neurobiological mechanisms *in vivo*, measuring 5,543 CSF metabolites, the largest panel in CSF to date, in 977 individuals of European ancestry. Individuals originated from two separate cohorts including cognitively healthy subjects (n=490) and a well-characterized memory clinic sample, the Amsterdam Dementia Cohort (ADC, n=487). We performed metabolite quantitative trait loci (mQTL) mapping on CSF metabolomics and found 126 significant mQTLs, representing 65 unique CSF metabolites across 51 independent loci. To better understand the role of CSF mQTLs in brain-related disorders, we performed a metabolome-wide association study (MWAS), identifying 40 associations between CSF metabolites and brain traits. Similarly, over 90% of significant mQTLs demonstrated colocalized associations with brain-specific gene expression, unveiling potential neurobiological pathways.

## Introduction

Genome-wide association studies (GWAS) have successfully identified many genetic risk loci contributing to human diseases and traits. The functional interpretation of these risk loci is crucial for treatment development and biomarker identification, but has been challenging. Quantitative trait loci (QTL) studies provide valuable mechanistic insights for various disease etiologies.^1^ The study of metabolomics in particular allows the detection of small changes in endogenous and exogenous compounds, thereby closely reflecting the current physiological state of cells, tissue or organisms.^2–6^

Due to the relative inaccessibility of *in vivo* brain tissue and its surroundings, (early) insights into biological processes underlying psychiatric and neurodegenerative disorders have been limited.^7,8^ Cerebrospinal fluid (CSF) is in close contact with the brain, and is important for providing nutrients and participates in waste removal of neural metabolism through interaction with interstitial fluid surrounding brain cells.^9^ CSF can be collected safely in humans through routine procedures, and thus provides an unique opportunity to study ongoing metabolic processes in the brain. CSF studies in neurodegenerative disease have been successful in identifying biological processes involved, such as abnormal regulation of lipid metabolism and increased inflammation.^10–12^ So far, CSF QTL mapping studies have shown that the metabolome^13–15^ is (at least partly) under genetic control. This implies that insight in genetic architecture of CSF metabolites can facilitate the functional understanding of genetic risk loci for brain-related traits.

In this study, we performed a genome-wide metabolite-QTL (mQTL) study on the largest mass-spectrometry (MS)-based CSF metabolomic panel studied so far (5,543 metabolites) using data from 977 individuals of European ancestry, including a cohort of cognitively healthy subjects (n=490) and a well-characterized memory clinic sample that is part of the Amsterdam Dementia Cohort (ADC, n=487). The assayed metabolites include those of primary metabolism, biogenic amines, and complex lipids. We identified 82 significant mQTLs for 65 CSF metabolites and 51 independent loci, of which 58/65 (89.2%) have not been detected before in CSF and eight not in blood, saliva or urine. This study indicates that the multi-omic integration of genetics with CSF metabolomic data helps to identify molecular neurobiological mechanisms.

## Results

### QTL mapping of CSF metabolites

Demographic and clinical characteristics are presented in Table 1. CSF samples were assessed for metabolic compounds via three platforms that each measure different components of the metabolome: GC-TOF MS/MS (primary metabolism, 393 metabolites, 273 of which are unannotated), CSH-QTOF MS/MS (complex lipids, 3,532 metabolites, 3,262 unannotated), and HILIC-QTOF MS/MS (biogenic amines, 1,618 metabolites, 1,194 unannotated) (Table S1). We did not observe genomic inflation for any of the CSF metabolites (range λ=0.97-1.03) (Table S1). Heritability estimates for CSF metabolite levels ranged from *h*^2^_SNP_=3%-49% (mean = 15%, SD = 12%), with the highest reported heritability for N-Acetylhistidine (49%), N-epsilon-Dimethyl-L-lysine (42%) and ethylmalonic acid (23%). Given a high degree of concordance between observed mQTL effects in both the cognitively healthy subjects cohort as the memory clinic cohort (Table S2), we performed a meta-analysis in the combined dataset of 977 subjects to boost power to detect associations (Figure 1).

**Figure 1.**
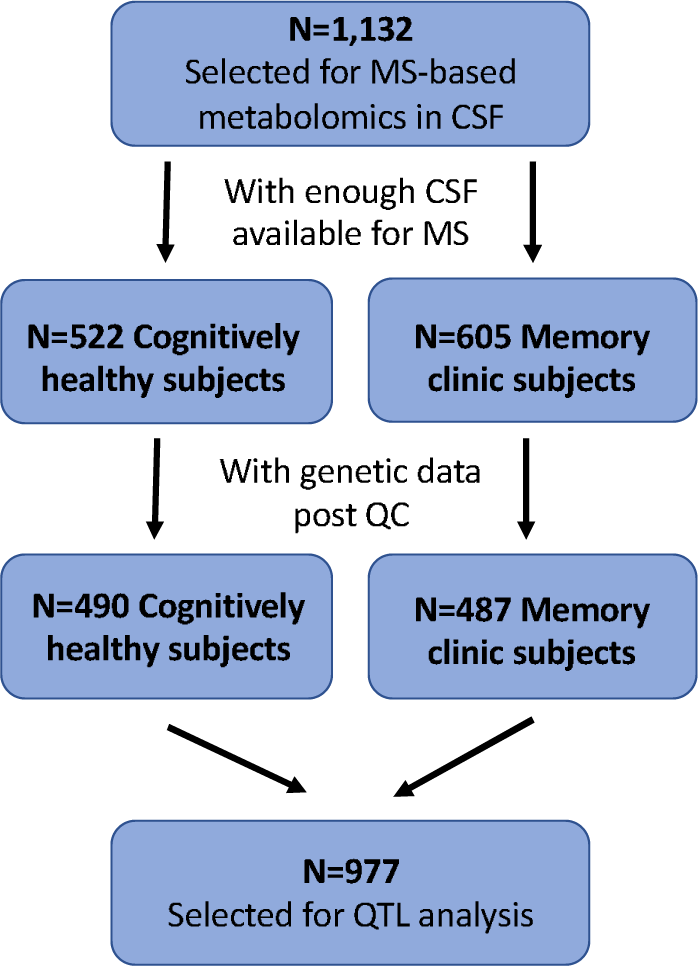
Overview of samples included in the CSF QTL mapping study.

**Table 1.** Demographics study samples.

|  | Cognitively healthy | Memory clinic |  | Memory clinic |  |  |  |
| --- | --- | --- | --- | --- | --- | --- | --- |
|  | all (N=490) | all (N=487) | p-value | Cognitively unimpaired subjects (n= 220, 45%) | Mild cognitive impairment (n= 87, 18%) | Dementia (n= 180, 37%) | p-value |
| Age, mean (SD) | 40.6 (10.8) | 65.5 (9.7) | 5,88E-188 | 65.2 (12) | 66.6 (7.7) | 65.3 (7.2) | 5,09E-01 |
| Female, n (%) | 144 (29%) | 207 (42.5%) | 1,90E-05 | 127 (58%) | 57 (66%) | 96 (53%) | 1,68E-01 |
| APOE-e4 at least one allele, n (%) | 172 (35%) | 246 (51%) | <0,00001 | 73 (33%) | 63 (72%) | 110 (61%) | 4,27E-12 |
| APOE-e2 at least one allele, n (%) | 75 (15.3%) | 34 (7%) | 5,60E-05 | 24 (11%) | 2 (2%) | 8 (4%) | 6,92E-03 |
| Abnormal Amyloid-beta, n(%) | NA | 329 (68%) | NA | 63 (29%) | 87 (100%) | 179 (100%) | 1,29E-61 |
| Abnormal p-tau, n (%) | NA | 235 (48%) | NA | 35 (16%) | 56 (64%) | 144 (80%) | 7,80E-39 |

### Detection of independent and novel CSF mQTL signals

Our genome-wide mQTL study on 5,543 metabolites and 6,189,630 SNPs identified 4,999 significant SNP-metabolite pairs after Bonferroni-correction (*P*<6e10^-11^). After applying a filter on independent LD blocks (based on clumping SNPs within 1Mb and LD R^2^<0.2) we observed 126 significant SNP-metabolite pairs, representing 65 unique CSF metabolite levels across 73 loci (Figure 2A). Following stepwise conditional analyses on the subset of CSF metabolites (n=19) with multiple independent loci (n=80), we identified in total 82 mQTLs (for 51 independent loci and 65 metabolites): 53 metabolites had one independent locus, eight metabolites had two loci, and four metabolites had three (or more) independent loci (Table 2, Table S3, Figure 2C). This is in line with previous studies, demonstrating that most of the heritability of metabolite levels is confined to a single locus,^13^ often uniquely regulated by the gene that encodes a protein involved in the specific metabolic pathway. CSF metabolites under a more complex regulatory architecture included ribonic acid (*ENOSF1/TYMS*), ethylmalonic acid (*UNC119B*), and two unannotated metabolites (i.e., *PYROXD2* for 9.31_161.13, *TYMS* for 9.40_165.04). Similarly, many independent mQTLs (n=33) harbored associations for more than one metabolite, involving a total of 39 different metabolites with at most five metabolites related to one locus (Figure 2D) (i.e., *FADS2*). This suggests pleiotropy at these loci, potentially involving a key enzyme in multiple metabolic pathways, or a group of metabolites that are all part of a single pathway.

**Figure 2.**
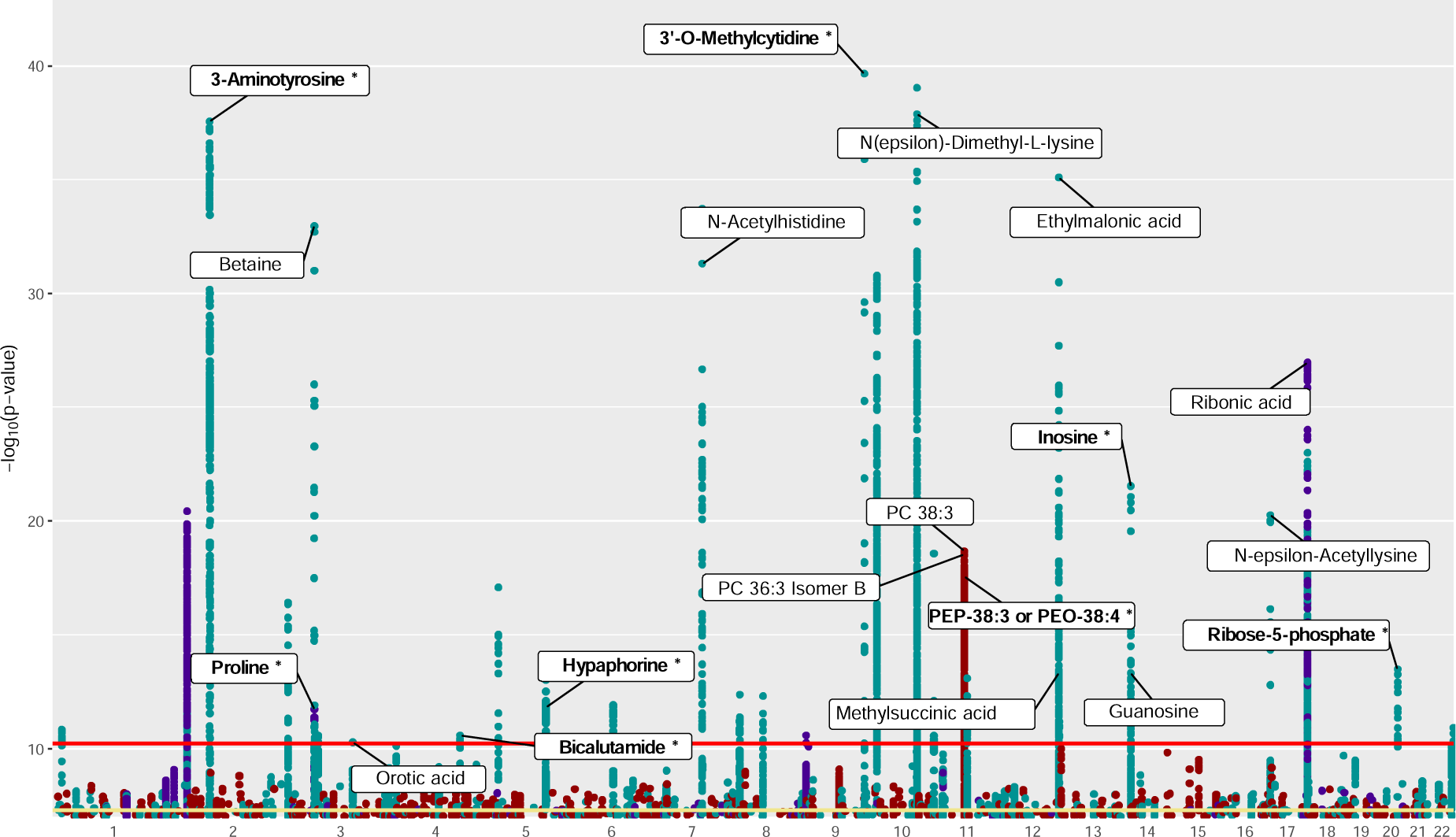
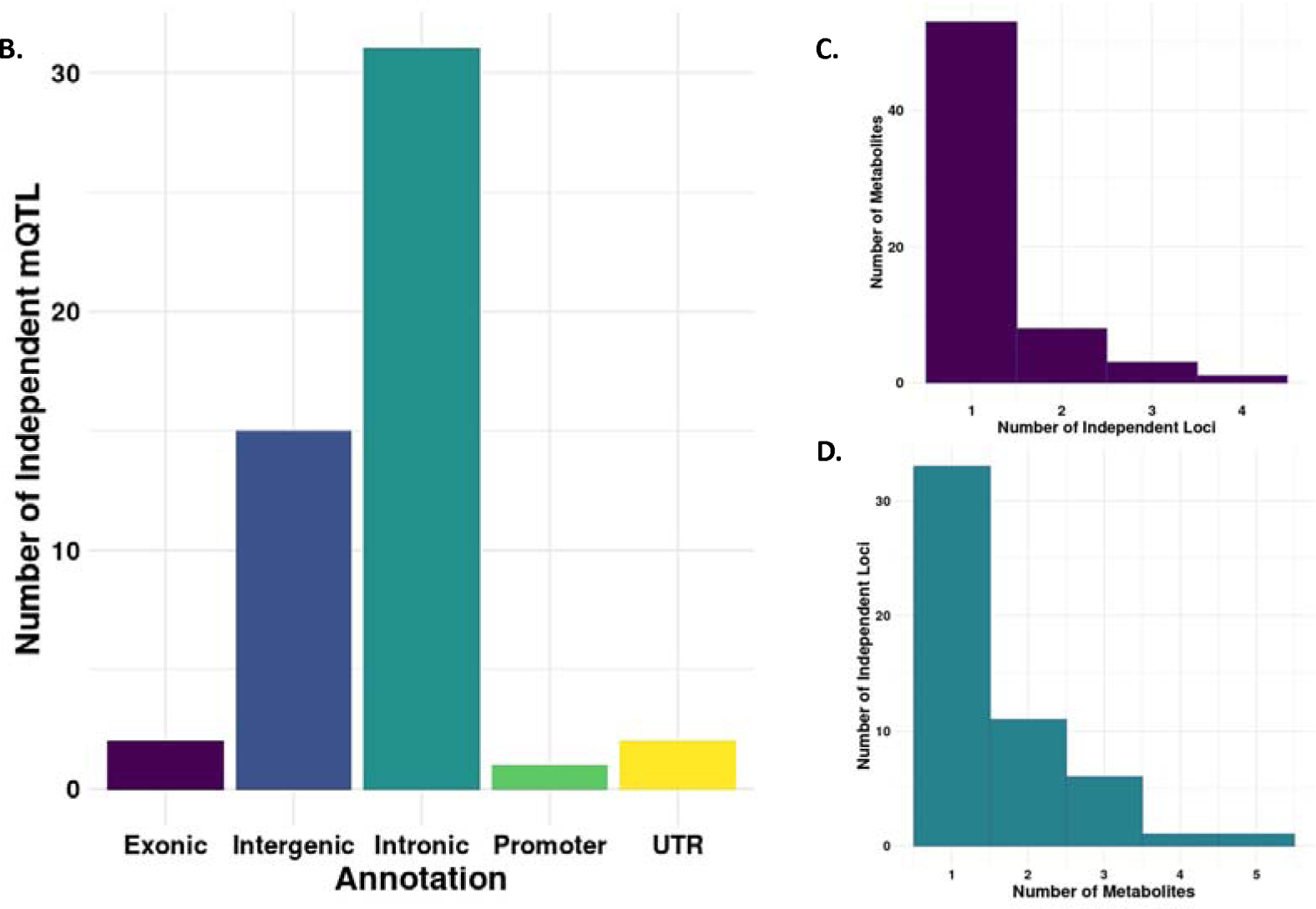
Manhattan plot and genetic architecture of mQTL associations. **(A)** Manhattan plot of the CSF metabolites that had at least one genome-wide significant mQTL association (*P*<6e-11) in the total sample (n=977). Each point depicts a distinct mQTL association between genetic variant and CSF metabolite levels. Green represents CSF metabolites measured using HILIC-QTOF MS/MS (biogenic amines, 1,618 metabolites), blue represents metabolites measured with GC-TOF MS (primary metabolism, 393 metabolites) and red represents CSH-QTOF MS/MS (complex lipids, 3,532 metabolites). Only CSF metabolites with annotation and *P*<6e-11 have a label. Novel CSF mQTL are depicted in bold and with an asterisk. The red line depicts the significance threshold of *P*<6e-11 (Bonferroni correction for 754 independent CSF signals), the yellow line depicts *P*<5e-8. Only results with a *P*<1e-8 and *P*>1e-40 are plotted.

**(B)** Using the GENCODE basic gene annotation, we first subsetted to 19,100 protein-coding genes, and we counted exons and UTRs as defined within the file. Promoters were defined as the region 10Kb upstream of a gene TSS, introns as regions within gene annotations not already covered by exons, and intergenic regions as any region not covered by any previous annotations. The y-axis denotes the number of independent SNPs, with some SNPs counted more than once for multiple metabolites.

**(C)** Histogram of polygenicity depicting number of metabolites associated with each independent locus.

**(D)** Histogram of pleiotropy depicting the number of independent loci associated with each metabolite.

**Table 2.** Overview of 126 significant mQTLs, representing 65 unique CSF metabolite levels across 51 independent loci.

| Nearest gene | LeadSNP | Metabolite | CHR | BP | Beta | P | mQTL<br>observed<br>before in |
| --- | --- | --- | --- | --- | --- | --- | --- |
| *AKR7A2* | chr1:19299791:G:T | 8.19_138.97 | 1 | 19299791 | 0,5045 | 2,92E-11 | N |
| *AKR7A2* | chr1:19299791:G:T | 8.19_94.98 | 1 | 19299791 | 0,5078 | 1,47E-11 | N |
| *UGT1A1* | rs1976391 | 1.05_584.26 | 2 | 233757337 | 0,3321 | 7,19E-12 | N |
| *UGT1A1* | rs1976391 | 1.04_583.26 | 2 | 233757337 | 0,4048 | 3,83E-17 | N |
| *ALMS1P1* | rs60762909 | 3-Aminotyrosine | 2 | 73646375 | -0,8882 | 6,96E-67 | N |
| *KHK* | rs6754371 | 100841 | 2 | 27092196 | 0,355 | 2,69E-15 | N |
| *KHK* | rs6754371 | 100343 | 2 | 27092196 | 0,4268 | 3,73E-21 | N |
| *EMILIN1* | rs7602823 | 8.12_275.05 | 2 | 27086538 | 0,3187 | 1,43E-12 | N |
| *TKT* | chr3:53245509:C:T | 8.40_275.05 | 3 | 53245509 | -0,3039 | 2,64E-11 | N |
| *SLC6A20* | rs17279437 | Betaine | 3 | 45772602 | 0,9191 | 1,08E-33 | CSF |
| *ITGB5* | rs2055983 | Oroticacid | 3 | 124761583 | -0,373 | 5,28E-11 | blood |
| *SACM1L* | rs73058498 | proline | 3 | 45723640 | 0,5816 | 1,88E-12 | N |
| *POU4F2* | rs1155168 | Bicalutamide | 4 | 146672579 | 0,425 | 2,63E-11 | N |
| *SLC22A5* | chr5:132389258:T:C | Hypaphorine | 5 | 132389258 | -0,4348 | 1,52E-12 | N |
| *SLC22A5* | chr5:132389258:T:C | 5.58_305.20 | 5 | 132389258 | -0,4381 | 1,15E-12 | N |
| *SLC22A5* | chr5:132389258:T:C | 5.58_188.07 | 5 | 132389258 | -0,4327 | 1,79E-12 | N |
| *AGXT2* | rs37369 | 6.21_174.12 | 5 | 35037010 | -0,5958 | 8,32E-18 | N |
| *SLC22A4* | rs56399423 | 6.02_160.13 | 5 | 132336964 | 0,3148 | 9,68E-14 | N |
| *PM20D2* | rs1048641 | 8.37_247.13 | 6 | 89165138 | -0,3497 | 1,24E-12 | N |
| *MOGAT3* | chr7:101200730:C:T | 7.99_210.09 | 7 | 101200730 | -0,7965 | 6,34E-23 | N |
| *MOGAT3* | chr7:101200730:C:T | 8.07_212.10 | 7 | 101200730 | -0,8072 | 1,99E-23 | N |
| *NT16* | rs740104 | N-Acetylhistidine | 7 | 101173066 | -1,36 | 8,33E-74 | CSF |
| *NT16* | rs740104 | 8.10_220.07 | 7 | 101173066 | -1,252 | 4,47E-61 | N |
| *NT16* | rs740104 | 7.40_268.13 | 7 | 101173066 | -0,7699 | 4,68E-25 | N |
| *NT2* | rs4921913 | 2.77_165.04 | 8 | 18414867 | -0,3528 | 1,63E-11 | N |
| *NT2* | rs4921913 | 5.42_183.05 | 8 | 18414867 | -0,3849 | 4,27E-13 | N |
| *NT2* | rs4921913 | 5.91_181.04 | 8 | 18414867 | -0,3339 | 4,33E-11 | N |
| *ADHFE1* | rs6993131 | 2.39_137.02 | 8 | 66449856 | -0,3459 | 4,92E-13 | N |
| *PHYHD1* | rs12005404 | 3'-O-Methylcytidine | 9 | 128935347 | 0,7544 | 9,48E-70 | N |
| *PTPRD* | rs679640 | 47 | 9 | 9581738 | 0,3149 | 2,67E-11 | N |
| *PTER* | chr10:16435878:G:C | 3.00_130.05 | 10 | 16435878 | 0,4449 | 1,97E-22 | N |
| *PTER* | chr10:16465256:CTAAGG:C | 2.92_130.05 | 10 | 16465256 | -0,4817 | 5,13E-27 | N |
| *PYROXD2* | chr10:98392298:C:G | 9.29_260.20 | 10 | 98392298 | -1,098 | 5,03E-181 | N |
| *PYROXD2* | chr10:98392298:C:G | 7.25_144.10 | 10 | 98392298 | -0,7816 | 2,20E-82 | N |
| *PYROXD2* | rs10883083 | N.epsilon.-Dimethyl-L-lysine | 10 | 98386156 | -1,112 | 1,80E-185 | CSF |
| *PYROXD2* | rs10883083 | 9.30_130.09 | 10 | 98386156 | -1,147 | 1,42E-203 | N |
| *PYROXD2* | rs10883083 | 9.31_161.13 | 10 | 98386156 | -1,161 | 3,58E-214 | N |
| *PTER* | rs2010442 | 3.04_132.06 | 10 | 16442889 | -0,5215 | 1,63E-31 | N |
| *PAOX* | rs4838735 | 6.62_230.19 | 10 | 133382797 | 0,4471 | 2,69E-19 | N |
| *GSTP1* | rs1695 | 9.00_365.17 | 11 | 67585218 | -0,3457 | 4,89E-13 | N |
| *GSTP1* | rs1695 | 9.29_422.20 | 11 | 67585218 | -0,3528 | 8,16E-14 | N |
| *FADS2* | rs174556 | PC38:3 (isomer 1) | 11 | 61813163 | 0,417 | 1,89E-18 | blood |
| *FADS2* | rs174556 | PC36:3IsomerB (isomer 1) | 11 | 61813163 | 0,4286 | 2,11E-19 | blood |
| *FADS2* | rs174556 | PC36:3IsomerB (isomer 2) | 11 | 61813163 | 0,3691 | 8,52E-15 | blood |
| *FADS2* | rs174556 | PC38:3 (isomer 2) | 11 | 61813163 | 0,4273 | 2,94E-19 | Y |
| *FADS2* | rs174559 | PEP-38:3orPEO-38:4 | 11 | 61814184 | 0,4231 | 2,71E-18 | N |
| *UNC119B* | rs12829722 | Ethylmalonicacid | 12 | 120717819 | 0,7332 | 1,79E-49 | CSF, blood,<br>saliva,<br>urine |
| *UNC119B* | rs12829722 | Methylsuccinicacid | 12 | 120717819 | 0,3876 | 4,97E-14 | blood,<br>saliva,<br>urine |
| *PNP* | chr14:20471430:G:T | Guanosine | 14 | 20471430 | 0,391 | 4,87E-14 | CSF |
| *PNP* | rs1713426 | 6.32_327.14 | 14 | 20467918 | 0,4026 | 8,07E-17 | N |
| *PNP* | rs1713426 | 6.32_269.09 | 14 | 20467918 | 0,3915 | 1,10E-15 | N |
| *PNP* | rs1713426 | 6.32_137.05 | 14 | 20467918 | 0,4095 | 7,45E-17 | N |
| *PNP* | rs1713427 | 6.32_313.12 | 14 | 20467999 | 0,361 | 2,36E-11 | N |
| *PNP* | rs1760940 | Inosine | 14 | 20470092 | 0,5151 | 2,91E-22 | N |
| *SAT2* | rs9913778 | N-epsilon-Acetyllysine | 17 | 7630583 | -0,771 | 5,46E-21 | Y |
| *TYMS* | rs10502289 | 9.82_165.04 | 18 | 676789 | -0,5221 | 1,90E-20 | N |
| *TYMS* | rs11081266 | 9.33_165.04 | 18 | 683441 | -0,47 | 4,04E-17 | N |
| *TYMS* | rs11081266 | 9.49_165.04 | 18 | 683441 | -0,4958 | 5,54E-19 | N |
| *TYMS* | rs11081266 | 9.23_165.04 | 18 | 683441 | -0,4831 | 7,68E-19 | N |
| *TYMS* | rs11081266 | 9.67_165.04 | 18 | 683441 | -0,4908 | 2,97E-18 | N |
| *TYMS* | rs11081266 | 9.40_165.04 | 18 | 683441 | -0,5446 | 1,01E-23 | N |
| *TYMS* | rs2790 | 9.22_309.06 | 18 | 673086 | -0,422 | 4,63E-14 | N |
| *ENOSF1/TYMS* | rs2847334 | ribonicacid | 18 | 692095 | 0,447 | 1,09E-27 | blood,<br>saliva,<br>urine |
| *SLC13A3* | rs381118 | Ribose-5-phosphate | 20 | 46574387 | -0,4931 | 3,24E-14 | N |
| *KLHDC7B* | rs131784 | 4.41_293.05 | 22 | 50543007 | 0,3501 | 1,23E-11 | N |

Most of the 82 mQTL associations were observed from the HILIC-QTOF MS/MS platform (69, 84.1%, SNP-metabolite associations), followed by GC-TOF (8, 9.8%, SNP-metabolite associations) and CSH-QTOF (5, 6.1%, SNP-metabolite associations).

### Most CSF mQTLs located in intronic regions

We examined the functional annotation of our identified CSF mQTLs and found that variants within intronic regions of protein coding genes account for 60.7% of loci, variants within promoters (defined as 10kb upstream of transcription start site) of genes account for 1.9% of independent loci, variants within untranslated regions (UTRs) for 3.9%, and variants within exons for 3.9% (Figure 2B). The paucity of exonic variants is a known trend among QTL and GWAS studies,^16,14,17^ whereas the abundance of intronic variants suggests potential role of enhancer activity^18,19^ in the regulation of these metabolites. The remaining variants (29.4%) were labeled as intergenic, for which functional annotations are lacking.

### QTL mapping of CSF metabolites identifies novel and validates known biological pathways

Of the twenty mQTL-associated CSF metabolites that were mapped to known metabolite names, thirteen (65%) have been previously detected in QTL studies on CSF^13^ (n=5), blood^2–4^ (n=10), saliva^5^ (n=3) or urine^6^ (n=3) metabolite levels (Figure S1, matched by metabolite then for SNPs in LD R2>0.8).

Eight annotated CSF mQTLs were novel (not previously reported in CSF, blood, saliva or urine), including those for hypaphorine, bicalutamide, 3’-O-Methylcytidine, PEP-38:3 or PEO-38:4, inosine, ribose-5-phosphate, 3-Aminotyrosine and proline (Table S4). One novel mQTL consisted of SNPs within the *SLC22A5* locus and CSF levels of exogenous metabolite hypaphorine (chr5:132389258:T:C, *P*=1.52e-12, no causal SNPs observed with fine-mapping) (Figure S10-S11). Another novel mQTL locus included the *SACM1L* locus on chromosome 3 and associated with CSF proline levels (rs73058498, *P*=1.88e-12). Statistical and functional fine-mapping with SuSiE selected the lead SNP rs73058498 (posterior inclusion probabilities (PIPs) range 0.25 - 0.40) as part of the only 95% credible set for proline (see Table S9 for fine-mapping SNPs within 95% credible sets for which their summed PIPs exceed 50%).

The CSH-QTOF platform for complex lipids harbored one mQTL locus, representing both previously identified and novel *pleiotropic* effects of *FADS2* (on chromosome 11) with CSF levels of four different phosphatidylcholines, PC 36:3 Isomer B (i.e., two isomers with different m/z ratios), PC 38:3 (two isomers) and and glycerophospholipid PE P-38:3/PE O-38:4 (novel association; rs174556, range *P*=8.52e-15 - 2.11e-19). Statistical and functional fine-mapping with FINEMAP highlighted the lead SNP rs174556 as part of a 50% credible set (along with eight other SNPs) for three out of four of the lipids, and at least two methods selected rs174556 as part of 95% credible sets (along with six other SNPs) for two of those three lipids (Table S9).

The strongest association was reported for *PYROXD2* on chromosome 10 with CSF levels of five inter-correlated metabolites (Pearson’s R2 > 0.83): N(epsilon)-Dimethyl-L-lysine and unannotated CSF metabolites 7.25_144.10, 9.29_260.20, 9.31_161.13, 9.30_130.09 (top SNP rs10883083 and chr10:98392298, range *P*=5.03e-181 - 3.58e-214 for all CSF metabolites) (Figure S2 - S6). Fine-mapping prioritized six potentially causal SNPs, including top SNP rs10883083 (as well as rs10786415, rs2147896, rs4539242, rs59667296, rs942814) in a 95% credible set for N(epsilon)-Dimethyl-L-lysine across all four methods. The top SNP rs10883083 was also part of the 95% credible sets for the other unannotated CSF metabolites at this locus (Table S9). The *PYROXD2* locus has been shown to associate with CSF levels of a similar (but not identical) metabolite called N6-methyl-L-lysine, suggesting that this locus plays a role in the regulation of lysine metabolism.^13^

Another association was observed for *NAT16* (N-acetyltransferase 16) on chromosome 7 with CSF levels of three inter-correlated CSF metabolites, including N-Acetylhistidine and two metabolites without a mapped biological name/label: 8.10_220.07 and 7.40_268.13 (rs740104, *P*= 8.33e-74, *P*=4.47e-61, *P*=4.68e-25, respectively) (Figure S7-S9). Fine-mapping identified two 95% credible sets for N-Acetylhistidine and 8.10_220.07, one of which (size = 4) included the top SNP rs740104, the other included only two SNPs, rs12540617 (PIPs range 0.72-0.85) and rs2227653 (PIPs range 0.14 - 0.28). For 7.40_268.13 we found only one 95% credible set (size = 4), including the top SNP (Table S9). N-Acetylhistidine has been found to play a role in the brain of some vertebrae, however it is still uncharacterized in humans.^20^ This association has been reported before in blood plasma.^21^

Another previously detected mQTL locus (in human plasma^22^, saliva^5^ and urine^6^) included four independent SNPs on *ENOSF1* (on chromosome 18) that associated with CSF levels of ribonic acid (rs11081229, *P*=2.36e10^-12^, rs2790, *P*=1.49e10^-27^, rs2847334, *P*=1.09e^-27^, rs6506537, *P*=1.27e10^-20^). Fine-mapping revealed two 95% credible sets for the ribonic acid mQTL, though only the SuSiE method prioritized the top SNP rs2847334 as part of a 95% credible set. Both SuSiE and PolyFun-SuSiE highlighted rs11081266 (PIPs range 0.19 - 0.41) and rs2790 (PIPs range 0.25 - 0.37) as the SNPs with the highest PIPs in a 95% credible set (Table S9).

### CSF metabolite levels associate with risk loci for brain-related disorders

To gain insight into the relevance of the genetic architecture underlying CSF metabolome to brain-related disorders, we performed metabolome-wide association (MWAS) analyses using the framework provided by FUSION.^23^ Out of the 220 CSF metabolites that had sufficient predictability (i.e., at least one SNP with *P*<5e-8), the *R*^2^ between the predicted and actual metabolite levels ranged from 0.005 to 0.67 (mean[=[0.05, SD[=[0.09). The HILIC-QTOF platform had the best performance and included more CSF metabolites as compared to the other platforms (Figure S12-13, Table S5).

CSF levels of 220 metabolites were tested for association with ten brain-related phenotypes, including Alzheimer’s disease^24^, dementia with Lewy bodies^25^, stroke^26^, amyotrophic lateral sclerosis^27^, bipolar disorder^28^, schizophrenia^29^, major depressive disorder^30^, attention deficit hyperactivity disorder (ADHD)^31^, insomnia^32^, and alcohol abuse disorder.^33^ We identified 40 significant (FDR<0.05) metabolite-phenotype associations, including ten traits and 31 CSF metabolite levels (Figure 3, Figure S14, Table S6). The strongest MWAS associations (with annotation) included: bipolar disorder with CSF metabolite levels of PC 36:3 Isomer B, PC 38:3, PE P-38:3/P EO-38:4 (*FADS2* on chromosome 11, range *P-*FDR=5.9e-08 - 4.6e-09), schizophrenia with CSF metabolite levels of 3-Aminotyrosine (*EMX1* on chromosome 2, *P-* FDR=4.25e-05). Three metabolite-trait associations had supporting evidence from colocalization analysis (posterior probability (PP)4>0.8), including bipolar disorder-PC36:3 Isomer B (*FADS2*), bipolar disorder-aspartic acid (intergenic region on chromosome 18 most nearby gene *CDH2*, rs2847408) and schizophrenia-2.06_142.09 (*SETD7*). Of these metabolome-wide significant genes with supporting colocalization evidence, the association of the intergenic region nearby *CDH2* for bipolar disorder was novel, showing no genome-wide evidence for association in the corresponding GWAS (minimal *p* within ±1 Mb of the gene’s region=2.06e-05)^28^.

**Figure 3.**
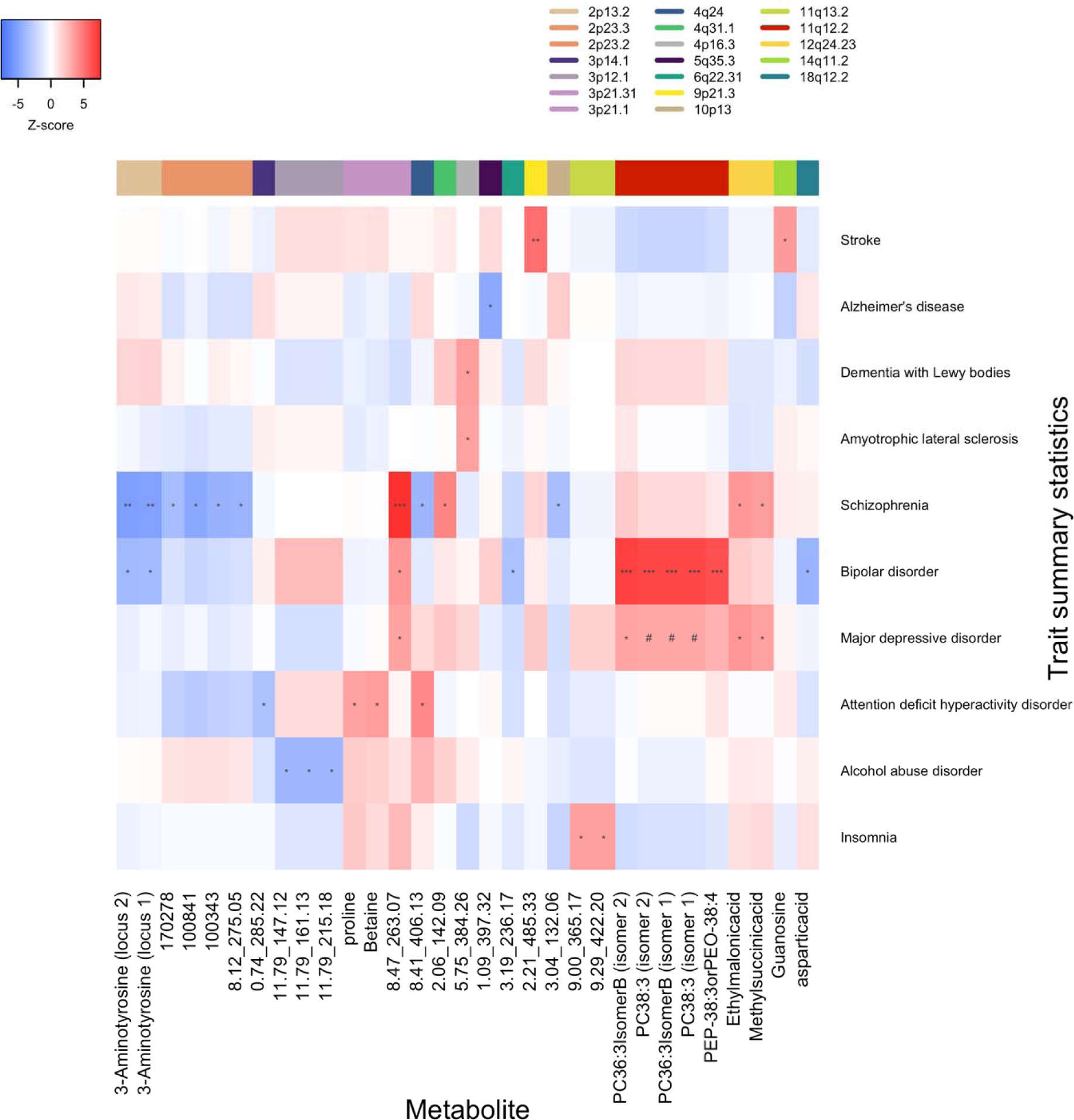
Heatmap of Z scores of CSF metabolites with at least one significant metabolome-wide significant association with one of the brain-related traits. This visualization includes metabolite-phenotype associations with at least one significant (*P*- FDR<0.05) association. Blank squares indicate that CSF metabolite weights were not sufficiently predictive. # depicts *P*-FDR<0.10, * *P*-FDR<0.05, ** *P*-FDR<0.0001*** and *P*-FDR<0.00001.

### Identified CSF mQTLs colocalize with known brain eQTLs

To identify whether any CSF mQTL colocalizes with eQTL for brain-specific gene expression, we used the PsychENCODE TWAS weights for 13,490 genes in a FUSION association test, followed by colocalization analysis. Of the 65 metabolites with a significant mQTL, we found that 37 (56.9%) have a high probability of colocalization (PP4>0.8) with brain-specific eQTL across 28 genes (Table S7).

We found that the mQTL on chromosome 14 regulating CSF levels of guanosine and inosine, both purine nucleosides, is highly likely to be colocalized (PP4>0.98) with the locus regulating expression levels of *PNP,* or purine nucleoside phosphorylase, in the brain. Within our dataset, guanosine and inosine levels are significantly correlated with an R2 of 0.65. Panyard, et. al previously reported the guanosine association at the *PNP* locus using CSF metabolomics, as well as an association with schizophrenia.^13^

Furthermore, we found a group of complex lipids that have high colocalization probability with a few genes at a locus on chromosome 11. Three of these lipids (two annotated as PC 36:3 Isomer B and PC 38:3) had colocalization PP4>0.9 with the *FADS1* (fatty acid desaturase 1) gene locus (Figure 4). These three lipids and an additional one (another PC 38:3 isomer) were colocalized (PP4>0.8) with the *FEN1* (flap endonuclease 1) gene locus. Similarly, these four lipids plus an additional one (annotated as “PE P-38:3 or PE O-38:4”) were colocalized (PP4>0.9) with the *TMEM258* (transmembrane protein 258) gene locus. Each of these lipids have a Pearson’s R2 > 0.83 with each other, suggesting that these lipids and their isomers are metabolites of a particular pathway. Sall, et. al.^34^ has previously grouped these genes together into a syntenic block of conserved MDD risk genes involved in myelination and lipid metabolism. The levels of these metabolites also showed highly significant associations with bipolar disorder in our MWAS.

**Figure 4.**
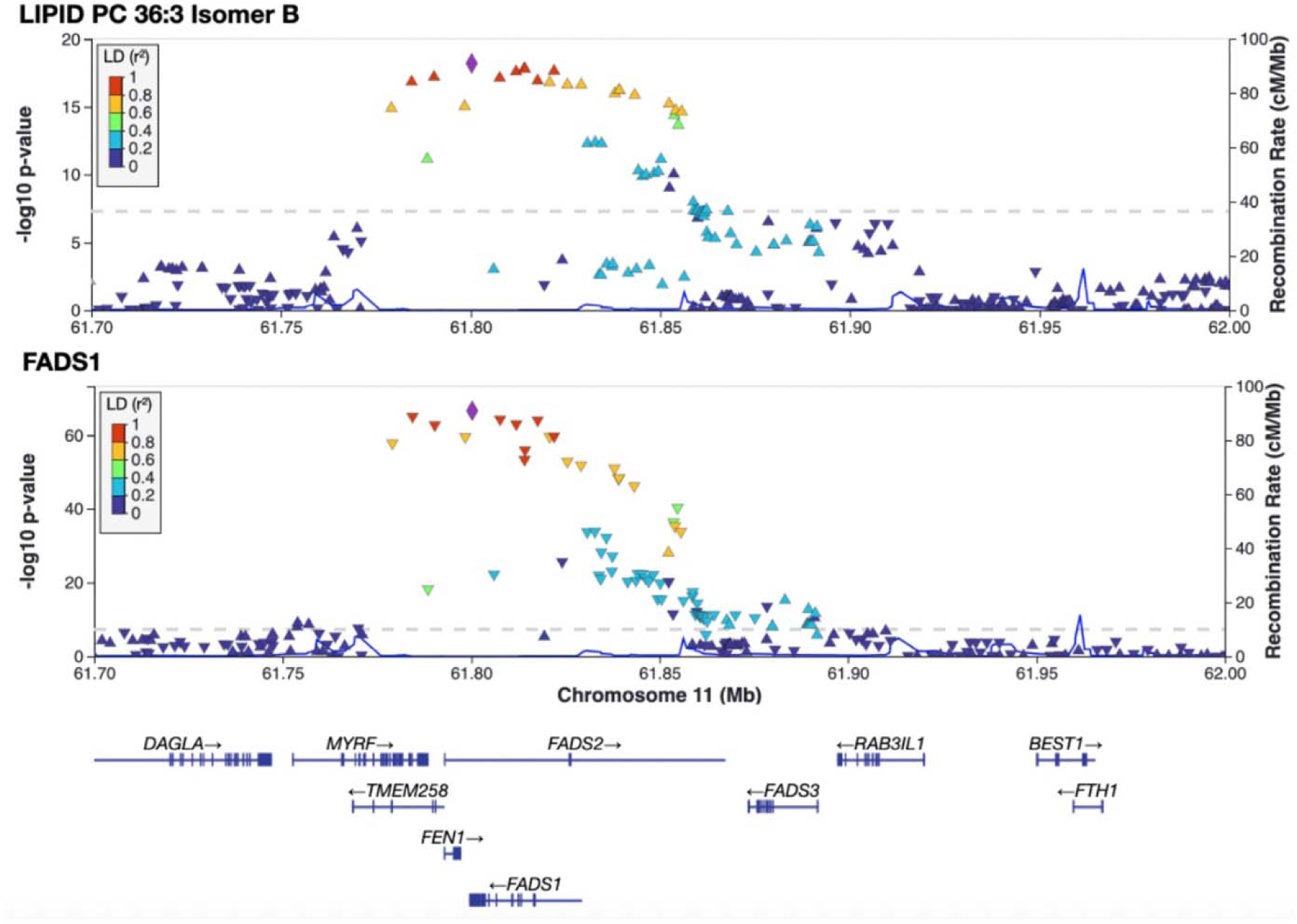
Phosphatidylcholine QTL colocalization with *FADS1* locus. Locus zoom plots, top panel shows associations with lipid PC 36:3 isomer B levels at the *FADS1*/*FADS2*/*TMEM258* locus and their LD relative to the top SNP; bottom panel shows eQTL SNPs for *FADS1*. The diamond shape denotes the lead SNP whereas the orientation of the triangles denote the direction of the effect. The gray dashed line indicates the significance threshold of p=5e-8.

Additionally, we applied the recently developed method isoTWAS^35^ in order to identify specific metabolite-associated isoforms that may otherwise be diluted in the gene-level TWAS approach. After adjusting p-values for multiple testing and applying a PIP threshold of 0.8, we find that 59 metabolite QTL (90.7%) are significantly associated across 96 different gene isoforms (61 unique genes) (Table S8). Of the 139 gene-metabolite pairs from isoTWAS, seven of these are direct replications of the TWAS findings (a significant overlap, Fisher’s exact test p-value = 3.6e-10), including the three of the lipids associated with the *FADS1* gene locus (as well as four unnamed HILIC metabolites associated with the genes *CTNNA1*, *PKD2L1*, *PYROXD2*, *SRSF12*). Many metabolites that were TWAS-significant were also isoTWAS-significant for different genes, including inosine and guanosine associated with isoforms of *EIF2B2* rather than *PNP*. EIF2B is known to have guanine nucleotide exchange factor activity,^36^ suggesting that isoTWAS is useful in identifying additional potential mechanisms of genetic regulation between metabolites and genes.

## Discussion

To gain more insight into the genetic architecture of the CSF metabolome, we performed genome-wide QTL mapping on 5,543 circulating CSF metabolites in a cohort of 977 living individuals. We identified 82 significant mQTLs representing 51 independent loci and 65 unique CSF metabolite levels, of which 58/65 (89.2%) have not been detected before in CSF and eight not in blood, saliva or urine.

We found both novel and replicated mQTL associations, of which many metabolites seemed to be regulated by the gene locus that encodes the protein for which they are substrates or products. One novel example found here is hypaphorine, a plant metabolite that can be found in the human metabolome after ingestion of legumes (e.g., lentils, chickpeas).^37^ Hypaphorine level was found to be regulated by the *SLC22A5* locus, a gene for the OCTN2 protein, which transports carnitine into the cell. Hypaphorine has been shown to have inhibitory effects on a SLC22 family members due to its similarity in structural backbone to carnitine.^38^ A strongly significant and replicated example is the association between ribonic acid and the *ENOSF1* locus, the protein for which catalyzes the dehydration of sugars, including ribonic acid.

Similarly, we observed that 56.9% (37/65) of the CSF metabolites with a mQTL colocalized with brain-specific eQTLs, and 90.7% (59/65) colocalized with brain specific isoform-QTLs (isoQTLs). Collectively, these eQTL-CSF mQTL colocalizations provide insights into the shared neurobiological mechanisms and possible interactions between these genes and CSF metabolite levels. One such eQTL-mQTL colocalization we found is at the *PNP* locus with CSF levels of guanosine and inosine. PNP is involved in the purine nucleotide salvage pathway by catalyzing the conversion of guanosine to guanine as well as inosine to hypoxanthine.

Our identified CSF mQTLs were integrated with pre-existing genome-wide summary statistics on ten psychiatric and neurodegenerative disorders, to improve our understanding of the relevance of CSF mQTLs in brain-related disorders. We identified 31 CSF metabolites of which predicted levels associated with brain-related traits. The strongest MWAS associations were with bipolar disorder and schizophrenia, of which an intergenic region nearby *CDH2* for bipolar disorder was novel. We also observed metabolite associations for Alzheimer’s disease, stroke, and ADHD, highlighting that the integration of genetic data with CSF metabolomics data helps to decipher biological pathways underlying brain-related disorders.

Several studies have previously reported the lipid mQTL hotspot at the *FADS1/2* locus,^39,13,14^ which we augment with additional lipids here. This genetic region has been described previously as both highly polygenic and pleiotropic, mostly influencing complex lipids synthesized from arachidonic acid, a product of the rate-limiting enzymes encoded by *FADS2* (and *FADS1*).^39^ This locus has also been identified as a GWAS risk region for bipolar disorder,^28,40^ and here we found this locus to be significantly associated with phosphatidylcholine lipid levels, glycerophospholipid levels, and bipolar disorder through our MWAS analysis. Colocalization with brain eQTLs and isoQTLs suggested a common genetic mechanism regulating the levels of these lipids and the levels of the *FADS1* gene. Heterozygous knockouts of the *FADS1/2* genes in mice have shown episodic phenotypes such as increases in hyperactivity (marked by increased wheel-running), depression-like episodes, and changes in circadian rhythms, suggestive of the symptoms commonly associated with bipolar disorder.^41^ Similarly, lipid dysregulation and genes involved herein are known to play a role in AD.^42^ While we did not find significant differences in the levels of these lipids in our ADC cohort (data not shown), genetic variants in the *FADS1* locus have also been implicated in increased risk for Alzheimer’s disease,^43^ suggesting further pleiotropy at this locus for this neurodegenerative phenotype as well.

This study successfully identified many mQTL associations in CSF that were replicated in two separately ascertained cohorts; nonetheless, several limitations should be taken into account. First, some metabolites had no annotations, or annotations with low reliability scores (based on the Metabolomics Standards Initiative (MSI) scale), which makes these associations difficult to interpret biologically. As an example, we observed a novel mQTL association for variants located on *POU4F2* and CSF levels of a metabolite annotated as a prostate cancer medicine, bicalutamide (MSI level=3, i.e., moderate reliability). However, we did not observe sex differences for this association despite bicalutamide being prescribed mainly in men for treating metastatic prostate cancer (bicalutamide inhibits testosterone by binding to androgen receptors). This suggests the potential that the identified metabolite represents another metabolite than bicalutamide. The POU4F2 protein (encoded by *POU4F2*) expression levels have been implicated in promoting tumor growth in various cancer types through the Hedgehog signaling pathway.^44^ Altogether this suggests that our newly identified mQTL could represent a true biological pathway for prostate cancer, but this could not be verified with the current annotation information on this metabolite. Furthermore, our study consisted solely of European-ancestry individuals. To gain a better understanding of the biology between CSF metabolites and genetic variation in brain related disorders, as well as to improve the fine-mapping accuracy of these QTL, many more samples of diverse ancestries must be included in future studies.^45^

Since CSF collection can occur *in vivo*, our results highlight that the specific integration of genetics with CSF metabolomic data could help understanding how genetic factors contribute to ongoing molecular mechanisms underlying neuropsychiatric disorders and neurobiological mechanisms. This study indicates that the multi-omic integration of genetics with CSF metabolomic data helps to identify which molecular mechanisms underlie diseases and disorders of the brain. Large-scale genome-wide studies on the CSF metabolome are limited, thus expanding on these studies is imperative for gaining biological insight in psychiatric and neurodegenerative disorders. As the summary statistics generated by this study can also be useful for studying other diseases and traits, data will be shared with the scientific community via the European Bioinformatics Institute GWAS Catalog (GCP000726).

## Methods

### Study sample(s)

In total, 977 study samples (age 52.7±16.6 years, 35.9% female) were included from a memory clinic cohort and a cohort of cognitively healthy subjects in the Netherlands.

*Memory clinic cohort:* Memory clinic samples originated from three cohorts, that are all related to Alzheimer center Amsterdam, including the Amsterdam Dementia Cohort (ADC),^46^ the 90+ study^47^ and the Twin Study.^48,49^ The ADC started collecting samples in the year 2000 and is an ongoing, observational follow-up study of patients who visited the memory clinic of the Alzheimer Center at Amsterdam UMC, location VU University medical center (VUmc).^46^ Dementia was diagnosed according to diagnostic guidelines for neurodegenerative disease.^50–53^ The Innovative Medicine Initiative European Information Framework for AD (EMIF-AD) 90+ study includes cognitively healthy elderly above 90 years old, and is aimed to identify factors associated with resilience to cognitive impairment in the oldest-old.^47^ Monozygotic twins (one subject per twin pair) were invited from the Netherlands Twin Register^48^ to participate in the PreclinAD study as part of the EMIF-AD project (http://www.emif.eu/).^49^ A detailed description of the cohorts is provided in the supplemental materials.

*Cognitively healthy subject cohort:* Cognitively healthy volunteers were recruited at outpatient pre-operative screening services in four hospitals in Utrecht, the Netherlands (August 2008 until March 2010).^15^ We included patients undergoing spinal anesthesia for minor elective surgical procedures, who ranged between 18 and 60 years of age and had all four grandparents born in The Netherlands or other Northwestern European countries (Belgium, Germany, UK, France and Denmark). Each candidate participant received a telephone interview to exclude subjects with self-reported psychotic or major neurological disorders (e.g., stroke, brain tumors, neurodegenerative diseases) and to record any use of psychotropic medication.

An overview of characteristics from the cognitively healthy subjects cohort (N=490, all cognitive healthy controls) and memory clinic cohort (N=487) is presented in Table S10. The Amsterdam sample includes n=220 cognitively unimpaired subjects, n=87 subjects with mild cognitive impairment (MCI) and n=180 patients with dementia. Patient groups in the Amsterdam sample differed from each other as expected, with the AD-type dementia group including more *APOE-* ε4 carriers, less *APOE-*ε2 carriers and having more abnormal AD CSF biomarkers compared to the MCI and cognitively unimpaired subjects. Cognitively healthy subjects were less often female, were younger, had a lower *APOE-*ε4 and a higher *APOE-*ε2 frequency, as compared to the memory clinic subjects.

All participating studies were approved by their respective Medical Ethics Committee. Informed consent, either from the patient or from the legal representative, was obtained from all participants.

### CSF data collection

*Memory clinic cohort:* CSF samples were collected via lumbar puncture, using a 25-gauge needle and syringe. CSF levels of amyloid-beta 42 (Aβ42), total tau (t-tau) and hyperphosphorylated 181 tau (p-tau) were determined as part of the diagnostic work-up, using enzyme-linked immunosorbent assays (ELISA) (Innotest: Fujirebio, Ghent, Belgium).^46^ For the ADC cohort, CSF Aβ42 values were adjusted for drift over time as described previously.^54^ Biomarker abnormality cut-offs for the ADC cohort are CSF Aβ42 < 813 pg/ml, CSF t-tau > 375 pg/ml, ratio t-tau/ Aβ42 > 0.52.^55^

*Cognitively healthy subject cohort*: A sample of 6 ml of CSF was obtained from each subject via lumbar puncture and immediately stored in fractions of 0.5 and 1 ml at −80L°C, as described previously. ^15^

### CSF metabolite processing

In total, 5,543 CSF metabolites were measured across three different metabolite assays at the West Coast Metabolomics Center at UC Davis, including GC-TOF MS (primary metabolism), CSH-QTOF MS/MS (complex lipids), and HILIC-QTOF MS/MS (biogenic amines) (https://metabolomics.ucdavis.edu/).

Metabolite levels were first screened for missingness across each cohort, removing any metabolites which had missing data for >20% of individuals. For remaining metabolites, we imputed missing values to half the median value for the corresponding metabolite across the cohorts, reasoning that these metabolites are likely present in quantities too low to detect within these individuals, and as done previously^12^. Inverse-rank normalization was performed on all metabolites to ensure normality for QTL mapping.

### Genotyping and imputation

Memory clinic samples were genotyped with the Illumina Global Screening Array (GSA) and cognitively healthy control data was genotyped with the OmniExpress Exome array. Quality control prior to imputation has been described in depth elsewhere.^56^ Autosomal genotypes were first filtered for SNPs with <2% SNP-missingness and >5% minor allele frequency (MAF) using plink^57^ separately per cohort. Individuals with call rate <98% were excluded. Genotype vcf files were then uploaded into the TopMed server for imputation and liftover to hg38. Post-imputation quality was assessed by filtering variants with imputation R2>0.3, providing about 8 million SNPs per cohort for downstream analyses. Imputed genotypes were then merged between the two cohorts, and then once more filtered for variants with <2% SNP-missingness and >5% MAF.

### mQTL mapping pipeline

Plink v1.90b^57^ –linear function was used to perform the QTL mapping across each of the metabolites separately. Covariates were adjusted via the –covar flag, including age, sex, and first 3 genotyping PCs. Initially cohorts were analyzed separately and the resulting effect sizes for any metabolite-SNP pairs found to be genome-wide significant in either cohort were then checked for replication. For the Amsterdam cohort we performed QTL mapping with additional covariate adjustment for diagnostic status, which did not significantly change results (data not shown). The effect sizes of each of these loci per metabolite were tested for concordance via Pearson’s correlation. Meta analysis was performed after merging individual-level genotype data across both cohorts.

### Variant annotation

We downloaded the GENCODE v.41 basic gene annotation GTF file to interpret potential functional consequences of the loci found to be associated with metabolites. We manually define promoters as the region 10Kb upstream of the TSS (using bedtools^58^ –flank -s -r 10000 -l 0), introns as regions within “gene” annotations that are not already covered by “exons” (bedtools –subtract), and intergenic regions as any region not covered by any annotations (bedtools –complement). Independent loci were selected per metabolite by subsetting the genotype files to only the SNPs that reached at least p<6e10^-11^ significance, then clumping (using plink –clump command) these SNPs into separate regions with LD R2 < 0.2. All remaining SNPs for any metabolites were then concatenated, deduplicated, and reformatted into a BED format file for use with the bedtools –intersect command.

### CSF metabolome-wide association study pipeline

To identify metabolites whose local-regulated CSF level is associated with brain-related phenotypes, we performed MWAS analyses using FUSION software (http://gusevlab.org/projects/fusion). First, we generated weights for all CSF metabolites with at least one genome-wide significant signal in the cohorts combined (*P*<5e-8). We restricted to loci +/- 500Kb around the lead SNPs per each metabolite, with the note that some metabolites were polygenic and had more than one associated genetic loci. For polygenic metabolites, any significant SNPs outside of the +/-500Kb range of another genome-wide significant SNP were included as a separate model. Age, sex, study site and the first three genotype PCs were included as covariates in the –covar flag of the FUSION.compute_weights.R script.

We used the FUSION.assoc_test.R script (default settings) to test for association between the CSF metabolite weights and GWAS for Alzheimer’s disease^24^, dementia with Lewy bodies^25^, stroke^26^, amyotrophic lateral sclerosis^27^, bipolar disorder^28^, schizophrenia^29^, major depressive disorder^30^, ADHD^31^, insomnia^32^ and alcohol abuse disorder.^33^ To account for LD structure we used 1000 Genomes data (all ancestries, build 38) data as LD reference panel. The --coloc^59^ flag was included to perform colocalization on any metabolites that had an association with the trait of interest with *P*-TWAS < 0.05. Colocalization analysis further narrowed down the metabolite-trait associations to those with a single variant influencing both the CSF metabolite level and trait, and associations with a PP4>0.8 were classified as having supporting colocalization evidence.

Results on gene–tissue associations per phenotype were corrected for multiple comparisons using a 5% FDR significance threshold. Significant MWAS loci were identified as novel if the strongest associated SNP was not nominally significant (*P* > 1e-5) in the corresponding GWAS within ±1 Mb of the transcriptional start site of the gene’s region.

### Fine-mapping of mQTLs

To predict which SNP(s) within mQTL associations were most likely to be causal, we used FINEMAP,^60^ SuSiE,^61^ PolyFun-FINEMAP, and PolyFun-SuSiE^62^ as fine-mapping methods. The summary statistics for these metabolites were lifted over to hg19 to match the UKBB LD reference panel as well as the UKBB functional annotations^62^, both of these are composed of British ancestry individuals. Top loci to be fine-mapped were defined by selecting the SNP with the lowest p-value for each metabolite and including any SNPs within the region 500kb upstream and 500kb downstream of the lead SNP. Results were processed for credible sets with summed fine-mapping posterior inclusion probability (PIP) of at least 0.5 across fewer than 10 SNPs, and SNP-associations that overlapped across multiple methods were prioritized as most likely to be the causal.

### Integration with brain eQTLs

To identify mQTL that colocalize with eQTL for brain-specific gene expression, we lifted over the PsychENOCDE TWAS weights from hg19 to hg38 and then performed a FUSION TWAS analysis. Each of the summary statistics for metabolites with Bonferroni-significant loci were included as separate input GWAS for the FUSION.assoc_test.R script with the PyschENCODE TWAS weights, using the 1000 Genomes European-specific (EUR) LD panel. The --coloc 0.05 flag was included to perform colocalization on any metabolites that were associated with gene expression with *P*-TWAS < 0.05. We consider any metabolite-gene pairs with coloc.PP3 > coloc.PP4 and coloc.PP4 > 0.8 to be significantly colocalized.

In a similar manner, we also applied the isoform-level TWAS method^35^ using precomputed weights per isoform derived from adult PsychENCODE data, as provided by the isoTWAS developers. In total we tested 7,530 genes, each with varying numbers of isoforms, that had positive heritability with a p-value < 0.05 within the PsychENCODE data. Similar to the TWAS analyses, we used the 1000 Genomes EUR LD reference panel. We then performed probabilistic fine-mapping of the significant associations, and filtered the resulting statistics to those transcripts in the credible sets with an adjusted screen p-value less than 0.05, permutation p-value less than 0.05, and PIP >= 0.8.

## Data availability

GWAS summary statistics on all CSF metabolite levels that were generated as part of this study have been deposited to the European Bioinformatics Institute GWAS Catalog (https://www.ebi.ac.uk/gwas/) under accession no. GCSTXXXXX. GENCODE v.41 basic gene annotation GTF file from https://www.gencodegenes.org/human/stats_41.html UKBB LD reference panel available from Amazon S3 bucket s3://broad-alkesgroup-ukbb- ld/UKBB_LD/ UKBB functional annotations available from Amazon S3 bucket https://broad-alkesgroup-ukbb-ld.s3.amazonaws.com/UKBB_LD/baselineLF_v2.2.UKB.polyfun.tar.gz PsychENCODE TWAS weights are available from http://resource.psychencode.org/. PsychENCODE isoTWAS weights are available from https://zenodo.org/record/6795947#.Y8mi2-zMLBI.

Summary statistics on Alzheimer’s disease^24^, dementia with Lewy bodies^25^ and amyotrophic lateral sclerosis^27^ are publicly available at the European Bioinformatics Institute GWAS Catalog under accession no. GCST90027158, GCST90001390 and GCST90027163, respectively. Summary statistics on bipolar disorder^28^, schizophrenia^29^, major depressive disorder^30^, attention deficit hyperactivity disorder (ADHD)^31^ and alcohol abuse disorder^33^ are publicly available on the psychiatric genomics consortium (PGC) website (https://www.med.unc.edu/pgc/results-and-downloads).

The MEGASTROKE consortium, launched by the International Stroke Genetics Consortium, has published the stroke^26^ summary statistics at https://www.megastroke.org. Full insomnia^32^ summary statistics for UKB and the top 10,000 SNPs for 23andMe are available at https://ctg.cncr.nl/software/summary_statistics/.

## Supplemental figure and table captions

**Figure S1. Replication of our identified CSF mQTLs in previous studies on CSF, blood, urine and saliva.**

Of the 65 CSF metabolites with a CSF mQTL association, 20 metabolites had an annotation. For 13/20 out of the metabolites with annotation, the mQTL association has been reported previously.

**Figure S2. Relationship between rs10883083 on *PYROXD2* and CSF N(epsilon)-Dimethyl-L-lysine levels.**

In this visualization, groups are based on the carriership of the effect allele. For example, “AA” (purple) corresponds to subjects homozygous for the rs10883083-A allele, “GA” (brown) to heterozygous subjects and “GG” (pink) to subjects homozygous for the rs10883083-G allele.

Abbreviations: CSF: cerebrospinal fluid, *PYROXD2*: Pyridine Nucleotide-Disulphide Oxidoreductase Domain 2. Inverse-rank normalization was performed on all metabolites to ensure normality for QTL mapping.

**Figure S3. Relationship between chr10:98392298 on *PYROXD2* and CSF 7.25_144.10 levels.**

Abbreviations: CSF: cerebrospinal fluid, *PYROXD2*: Pyridine Nucleotide-Disulphide Oxidoreductase Domain 2. Inverse-rank normalization was performed on all metabolites to ensure normality for QTL mapping.

**Figure S4. Relationship between chr10:98392298 on *PYROXD2* and CSF 9.29_260.20 levels.**

Abbreviations: CSF: cerebrospinal fluid, *PYROXD2*: Pyridine Nucleotide-Disulphide Oxidoreductase Domain 2. Inverse-rank normalization was performed on all metabolites to ensure normality for QTL mapping.

**Figure S5. Relationship between chr10:98392298 on *PYROXD2* and CSF 9.30_130.09 levels.**

Abbreviations: CSF: cerebrospinal fluid, *PYROXD2*: Pyridine Nucleotide-Disulphide Oxidoreductase Domain 2. Inverse-rank normalization was performed on all metabolites to ensure normality for QTL mapping.

**Figure S6. Relationship between rs10883083 on *PYROXD2* and CSF 9.31_161.13 levels.**

Abbreviations: CSF: cerebrospinal fluid, *PYROXD2*: Pyridine Nucleotide-Disulphide Oxidoreductase Domain 2. Inverse-rank normalization was performed on all metabolites to ensure normality for QTL mapping.

**Figure S7. Relationship between rs740104 on *NAT16* and CSF N-Acetylhistidine levels.**

Abbreviations: CSF: cerebrospinal fluid, *NAT16*: N-acetyltransferase 16. Inverse-rank normalization was performed on all metabolites to ensure normality for QTL mapping.

**Figure S8. Relationship between rs740104 on *NAT16* and CSF 8.10_220.07 levels.**

Abbreviations: CSF: cerebrospinal fluid, *NAT16*: N-acetyltransferase 16. Inverse-rank normalization was performed on all metabolites to ensure normality for QTL mapping.

**Figure S9. Relationship between rs740104 on *NAT16* and CSF 7.40_268.13 levels.**

Abbreviations: CSF: cerebrospinal fluid, *NAT16*: N-acetyltransferase 16. Inverse-rank normalization was performed on all metabolites to ensure normality for QTL mapping.

**Figure S10. Manhattan plot of hypaphorine levels in CSF.**

Only associations with a *P* < 1.00 e-2 are plotted.

**Figure S11. Relationship between chr5:132389258:T:C on *SLC22A5* and CSF hypaphorine levels.**

Abbreviations: CSF: cerebrospinal fluid, *SLC22A5:* Solute Carrier Family 22 Member 5.

**Figure S12. Association (rsq) between actual and genetically predicted CSF metabolite levels according to the FUSION framework, stratified according to metabolite assays, including GC-TOF MS (primary metabolism), CSH-QTOF MS/MS (complex lipids), and HILIC-QTOF MS/MS (biogenic amines).**

Abbreviations: CSF: cerebrospinal fluid, GC-TOF MS: gas chromatography mass spectrometry, CSH-QTOF MS/MS: charged surface hybrid column quadrupole time-of-flight tandem mass spectrometry, HILIC-QTOF MS/MS: high pressure liquid chromatography quadrupole time-of-flight tandem mass spectrometry.

**Figure S13. Association (rsq) between actual and genetically predicted CSF metabolite levels according to the FUSION framework. Four different models were used to estimate predicted CSF metabolite levels. For each metabolite, the best performing model was selected in this TWAS analysis.**

Abbreviations: CSF: cerebrospinal fluid, blup: Best Linear Unbiased Predictor computed from all SNPs, enet: elastic-net regression, lasso: LASSO regression, top1: Single best QTL effect.

**Figure S14. Manhattan plot on metabolome-wide association results, representing the association between and CSF metabolite weights and GWAS for ten brain-related phenotypes.**

We identified 40 significant (FDR<0.05) metabolite-phenotype associations, including ten traits and 31 CSF metabolite levels. Each point depicts a distinct metabolite-trait association. MWAS hits with supporting evidence from colocalization analysis are highlighted in green. The red line depicts the significance threshold; pFDR < .05.

Abbreviations: CSF: cerebrospinal fluid, GWAS: genome-wide association study, FDR: false discovery rate; MWAS, metabolome-wide association study.

**Figure S15. Relationship between rs1155168 on *POU4F2* and CSF bicalutamide levels.**

Abbreviations: CSF: cerebrospinal fluid, *POU4F2*: POU Class 4 Homeobox 2

**Table S1. Feature characteristics of metabolites in cerebrospinal fluid (CSF) that passed the quality check. Lambda values represent the genomic inflation factors for each metabolite GWAS.**

**Table S2. Replication of mQTL between the memory clinic and cognitively healthy control cohorts.**

Memory clinic-specific (Amsterdam) and cognitively healthy control-specific p value and beta per each top SNP-metabolite pair are reported alongside meta-analysis p values and betas.

**Table S3. Results conditional analysis.**

Freq values represent the number of independent SNPs significantly associated with the metabolite before conditional analysis, and Freq.aftercond represents the number of independent SNPs still significant after conditional analysis.

**Table S4. Replication in other biofluids.**

**Table S5. Overview of number and proportion of CSF metabolites included in this study and included in the metabolome-wide association analysis.**

**Table S6. Metabolome-wide association study results on 10 brain-related disorders.**

Abbreviations: CHR; chromosome, P0; gene start, P1; gene end, HSQ; heritability of the gene, BEST.GWAS.ID; rsID of the most significant SNP in locus, BEST.GWAS.Z; Z-score of the most significant GWAS SNP in locus, EQTL.ID; rsID of the best eQTL in the locus, EQTL.R2; cross-validation R2 of the best eQTL in the locus, EQTL.Z; Z-score of the best eQTL in the locus, EQTL.GWAS.Z; GWAS Z-score for this eQTL, NSNP; number of SNPs in the locus, MODELCV.R2; cross-validation R2 of the best performing model, MODELCV.PV; cross-validation P-value of the best performing model, SNP; single nucleotide polymorphism, FDR; false-discovery rate, eQTL; expression quantitative trait locus.

**Table S7. Colocalized mQTL with psychENCODE eQTL.**

Abbreviations: CHR; chromosome, P0; gene start, P1; gene end, HSQ; heritability of the gene, BEST.GWAS.ID; rsID of the most significant SNP in locus, BEST.GWAS.Z; Z-score of the most significant GWAS SNP in locus, EQTL.ID; rsID of the best eQTL in the locus, EQTL.R2; cross-validation R2 of the best eQTL in the locus, EQTL.Z; Z-score of the best eQTL in the locus, EQTL.GWAS.Z; GWAS Z-score for this eQTL, NSNP; number of SNPs in the locus, MODELCV.R2; cross-validation R2 of the best performing model, MODELCV.PV; cross-validation P-value of the best performing model, SNP; single nucleotide polymorphism, FDR; false-discovery rate, eQTL; expression quantitative trait locus. PP; posterior probability (0: no association, 1: eQTL association only, 2: mQTL association only, 3: independent eQTL/mQTL association, 4: colocalized association).

**Table S8. isoTWAS results of significant metabolite-transcript pairs.**

Abbreviations: SNP; single nucleotide polymorphism, PIP; posterior inclusion probability; Z; Z-score of the association between the transcript and metabolite.

**Table S9. Fine-mapping 50% credible sets.**

The credible set column denotes which set the specific SNP was a part of specifically within each method. The locus column is defined as the top SNP for that metabolite, the SNP column is the fine-mapped SNP.

Abbreviations: CHR; chromosome, BP; base pair location, SNP; single nucleotide polymorphism, PIP; posterior inclusion probability, SD; standard deviation.

**Table S10. Demographics of the memory clinic (Amsterdam) and cognitively healthy control cohorts.**

## Supplemental methods

### Cohorts

Amsterdam Dementia Cohort (ADC): All patients receive a standardized multidisciplinary assessment as part of the diagnostic workup, consisting of medical history, informant-based history, neurological and medical examination, neuropsychological investigation, brain magnetic resonance imaging (MRI), standard laboratory work-up and lumbar puncture for cerebrospinal fluid (CSF) assessment.

The main aim of the EMIF-AD PreclinAD study was to identify new risk factors and diagnostic markers for both amyloid pathology and cognitive decline in cognitively normal subjects with or without amyloid pathology.

### MWAS colocalization analysis

MWAS colocalization analysis estimates the posterior probability (PP) for: 0. No association with either the brain-related trait or CSF metabolite levels (PP0), 1. Association with brain-related trait only (PP1), 2. Association with CSF metabolite levels only (PP2), 3. Association with brain-related trait and CSF metabolite levels from distinct causal variants (PP3) and 3. Shared causal variant (PP4). We defined colocalization evidence as supportive for a shared causal variant between the brain-related trait and CSF metabolite levels, when PP4>0.17 and PP4>PP3, as discussed in Gusev 2016.^23^

## Supplemental results

### Replication between cohorts

In order to quantify the concordance of the associations between both cohorts, for each metabolite, we subsetted the resulting summary statistics to any metabolite-SNP pairs associated with p<6e^-11^ in either cohort. In total, we identified 30 metabolites with significant genetic regulation in the cognitively healthy controls, and 29 metabolites in the memory clinic samples, of which 16 metabolites were significant after Bonferroni correction in both cohorts with the same direction of effect at the same SNPs. Of the remaining 27 mQTL that showed Bonferroni-significance in one cohort but not the other, 19 mQTL were nominally significant in the other cohort, with a p-value of at least 0.05 and same direction of effect at the same SNP, resulting in a direct replication rate of 81%. Out of the eight metabolite-SNP pairs that did not directly replicate, six were significant in the Memory clinic cohort but not replicated in the Cognitively healthy cohort, and the remaining two were not replicated in the Memory clinic cohort. No SNPs with LD R2 > 0.5 replicate with P < 0.05 in the opposing cohort, though when we consider the meta-analysis summary statistics, all SNP-metabolite associations are replicated with P <= 0.0088 (see Table S2). Furthermore, these eight specific instances each have at most two SNPs passing the Bonferroni-corrected threshold, suggesting that they are borderline significant associations.

### MWAS

Models include between 889 and 7431 SNPs. The top1 model was chosen as the best model type for 62.3% of the metabolites, followed by LASSO models (21.4%), elastic net models (10%), and blup (6.4%).

### Association for bicalutamide

We observed a novel mQTL association for variants located on *POU4F2* and CSF levels of the exogenous metabolite bicalutamide (rs1155168, *P*=2.63e-11) (Figure S15). Bicalutamide inhibits testosterone by binding to androgen receptors, and is prescribed for testosterone-related tumors as metastatic prostate cancer and to a lesser extent for PCOS symptoms (due to elevated levels of testosterone) in women. POU4F2 protein (encoded by *POU4F2*) expression levels has been shown to promote tumor growth in various cancer types through the Hedgehog signaling pathway^44^. Since this mQTL association comprises an exogenous metabolite, it may indicate that rs1155168-X associates with a higher risk for developing cancer (and thus using bicalutamide). We did not observe sex-differences for this association, while this could have been expected since bicalutamide is more often prescribed in men than women.

## Acknowledgements

We are greatly appreciative to those individuals who donated the CSF samples on which this study was based. We appreciate data acquisition and reporting by the UC Davis West Coast Metabolomics Center. We thank Dr. Evan Hurlow for his insightful discussions on the organic chemistry involved in this study. We would also like to thank Nicole Zelster for her contributions in the early stages of the project. We also thank Dr. Jurjen Luykx for his help with the recruitment of the cognitively healthy controls.

Research of Alzheimer Center Amsterdam is part of the neurodegeneration research program of Amsterdam Neuroscience. Alzheimer Center Amsterdam is supported by Stichting Alzheimer Nederland and Stichting VUmc Steun Alzheimercentrum Amsterdam. The clinical database structure was developed with funding from Stichting Dioraphte. This work has received support from the EU/EFPIA Innovative Medicines Initiative Joint Undertaking (EMIF grant number 115372) and Stichting Dioraphte. Genotyping of the Dutch case-control samples was performed in the context of EADB (European Alzheimer DNA biobank) funded by the JPco-fuND FP-829-029 (ZonMW projectnumber 733051061).

This project was funded by the NIH National Institute on Aging (NIA) grant RF1AG058484 and the National Institute of Mental Health (NIMH) grant R01MH115676 to RAO. Lianne Reus was funded by the Memorabel fellowship (ZonMW projectnumber: 10510022110012). Toni Boltz was supported by the NIH (grant no. 5T32HG002536).

S.L. is recipient of funding from ZonMW (#733050512), Health∼Holland, Topsector Life Sciences & Health (PPP-allowance; #LSHM20106). Stichting Dioraphte, the Edwin Bouw Fonds and Stichting VUmc fonds. W.vd.F., S.vd.L., C.T. are recipients of ABOARD, which is a public-private partnership receiving funding from ZonMW (#73305095007) and Health∼Holland, Topsector Life Sciences & Health (PPP-allowance; #LSHM20106). More than 30 partners participate in ABOARD (www.aboard-project.nl). ABOARD also receives funding from de Hersenstichting, Edwin Bouw Fonds and Gieskes-Strijbisfonds. The MEGASTROKE project received funding from sources specified at http://www.megastroke.org/acknowledgments.html.

## References

1. Kraus, W. E. et al. Metabolomic Quantitative Trait Loci (mQTL) Mapping Implicates the Ubiquitin Proteasome System in Cardiovascular Disease Pathogenesis. PLoS Genet. 11, e1005553 (2015).

2. Shin, S.-Y. et al. An atlas of genetic influences on human blood metabolites. Nat. Genet. 46, 543–550 (2014).

3. Prakash, S. Human metabolic individuality in biomedical and pharmaceutical research. Circulation. Cardiovascular genetics vol. 4 714–715 (2011).

4. Long, T. et al. Whole-genome sequencing identifies common-to-rare variants associated with human blood metabolites. Nat. Genet. 49, 568–578 (2017).

5. Nag, A. et al. Genome-wide scan identifies novel genetic loci regulating salivary metabolite levels. Hum. Mol. Genet. 29, 864–875 (2020).

6. Schlosser, P. et al. Genetic studies of urinary metabolites illuminate mechanisms of detoxification and excretion in humans. Nat. Genet. 52, 167–176 (2020).

7. Yang, C. et al. Genomic atlas of the proteome from brain, CSF and plasma prioritizes proteins implicated in neurological disorders. Nat. Neurosci. 24, 1302–1312 (2021).

8. Gusev, A. et al. Transcriptome-wide association study of schizophrenia and chromatin activity yields mechanistic disease insights. Nat. Genet. 50, 538–548 (2018).

9. Nedergaard, M. Garbage Truck of the Brain. Science vol. 340 1529–1530 Preprint at 10.1126/science.1240514 (2013).

10. Tijms, B. M. & Visser, P. J. Pathophysiological subtypes of Alzheimer’s disease based on cerebrospinal fluid proteomics. Alzheimer’s & Dementia vol. 16 Preprint at 10.1002/alz.037184 (2020).

11. Yan, J., Kuzhiumparambil, U., Bandodkar, S., Dale, R. C. & Fu, S. Cerebrospinal fluid metabolomics: detection of neuroinflammation in human central nervous system disease. Clin Transl Immunology 10, e1318 (2021).

12. Dong, R. et al. CSF metabolites associate with CSF tau and improve prediction of Alzheimer’s disease status. Alzheimers. Dement. 13, e12167 (2021).

13. Panyard, D. J. et al. Cerebrospinal fluid metabolomics identifies 19 brain-related phenotype associations. Preprint at 10.1101/2020.02.14.948398.

14. Tahir, U. A. et al. Whole Genome Association Study of the Plasma Metabolome Identifies Metabolites Linked to Cardiometabolic Disease in Black Individuals. Nat. Commun. 13, 4923 (2022).

15. Luykx, J. J. et al. Genome-wide association study of monoamine metabolite levels in human cerebrospinal fluid. Mol. Psychiatry 19, 228–234 (2014).

16. Watanabe, K. et al. A global overview of pleiotropy and genetic architecture in complex traits. Nat. Genet. 51, 1339–1348 (2019).

17. Flynn, E. D. & Lappalainen, T. Functional Characterization of Genetic Variant Effects on Expression. Annu Rev Biomed Data Sci 5, 119–139 (2022).

18. Jo, B.-S. & Choi, S. S. Introns: The Functional Benefits of Introns in Genomes. Genomics Inform. 13, 112 (2015).

19. Niu, H.-M. et al. Comprehensive functional annotation of susceptibility SNPs prioritized 10 genes for schizophrenia. Transl. Psychiatry 9, 1–12 (2019).

20. Baslow, M. H. & Guilfoyle, D. N. N-acetyl-l-histidine, a Prominent Biomolecule in Brain and Eye of Poikilothermic Vertebrates. Biomolecules 5, 635 (2015).

21. Yin, X. et al. Genome-wide association studies of metabolites in Finnish men identify disease-relevant loci. Nature Communications vol. 13 Preprint at 10.1038/s41467-022-29143-5 (2022).

22. Koshiba, S. et al. Identification of critical genetic variants associated with metabolic phenotypes of the Japanese population. Commun Biol 3, 662 (2020).

23. Gusev, A. et al. Integrative approaches for large-scale transcriptome-wide association studies. Nat. Genet. 48, 245–252 (2016).

24. Bellenguez, C. et al. New insights into the genetic etiology of Alzheimer’s disease and related dementias. Nat. Genet. 54, 412–436 (2022).

25. Chia, R. et al. Genome sequencing analysis identifies new loci associated with Lewy body dementia and provides insights into its genetic architecture. Nat. Genet. 53, 294–303 (2021).

26. Malik, R. et al. Multiancestry genome-wide association study of 520,000 subjects identifies 32 loci associated with stroke and stroke subtypes. Nat. Genet. 50, 524–537 (2018).

27. van Rheenen, W. et al. Common and rare variant association analyses in amyotrophic lateral sclerosis identify 15 risk loci with distinct genetic architectures and neuron-specific biology. Nat. Genet. 53, 1636–1648 (2021).

28. Mullins, N. et al. Genome-wide association study of more than 40,000 bipolar disorder cases provides new insights into the underlying biology. Nat. Genet. 53, 817–829 (2021).

29. Trubetskoy, V. et al. Mapping genomic loci implicates genes and synaptic biology in schizophrenia. Nature 604, 502–508 (2022).

30. Howard, D. M. et al. Genome-wide meta-analysis of depression identifies 102 independent variants and highlights the importance of the prefrontal brain regions. Nat. Neurosci. 22, 343–352 (2019).

31. Demontis, D. et al. Discovery of the first genome-wide significant risk loci for attention deficit/hyperactivity disorder. Nat. Genet. 51, 63–75 (2019).

32. Watanabe, K. et al. Genome-wide meta-analysis of insomnia prioritizes genes associated with metabolic and psychiatric pathways. Nat. Genet. 54, 1125–1132 (2022).

33. Sanchez-Roige, S. et al. Genome-Wide Association Study Meta-Analysis of the Alcohol Use Disorders Identification Test (AUDIT) in Two Population-Based Cohorts. Am. J. Psychiatry 176, 107–118 (2019).

34. Sall, S., Thompson, W., Santos, A. & Dwyer, D. S. Analysis of Major Depression Risk Genes Reveals Evolutionary Conservation, Shared Phenotypes, and Extensive Genetic Interactions. Front. Psychiatry 12, 698029 (2021).

35. Bhattacharya, A. et al. Isoform-level transcriptome-wide association uncovers extensive novel genetic risk mechanisms for neuropsychiatric disorders in the human brain. medRxiv 2022.08.23.22279134 (2022) doi:10.1101/2022.08.23.22279134.

36. Fogli, A. et al. Decreased guanine nucleotide exchange factor activity in eIF2B-mutated patients. Eur. J. Hum. Genet. 12, 561–566 (2004).

37. Keller, B. O., Wu, B. T. F., Li, S. S. J., Monga, V. & Innis, S. M. Hypaphorine is present in human milk in association with consumption of legumes. J. Agric. Food Chem. 61, 7654– 7660 (2013).

38. Yee, S. W. et al. Deorphaning a Solute Carrier 22 family member, SLC22A15, through functional genomic studies. FASEB J. 34, 15734 (2020).

39. Auwerx, C. et al. Exploiting the mediating role of the metabolome to unravel transcript-to-phenotype associations. Elife 12, (2023).

40. Ikeda, M. et al. A genome-wide association study identifies two novel susceptibility loci and trans population polygenicity associated with bipolar disorder. Mol. Psychiatry 23, 639–647 (2018).

41. Yamamoto, H. et al. GWAS-identified bipolar disorder risk allele in the FADS1/2 gene region links mood episodes and unsaturated fatty acid metabolism in mutant mice. Mol. Psychiatry (2023) doi:10.1038/s41380-023-01988-2.

42. Yin, F. Lipid metabolism and Alzheimer’s disease: clinical evidence, mechanistic link and therapeutic promise. FEBS J. 290, 1420–1453 (2023).

43. Hammouda, S. et al. Genetic variants in FADS1 and ELOVL2 increase level of arachidonic acid and the risk of Alzheimer’s disease in the Tunisian population. Prostaglandins Leukot. Essent. Fatty Acids 160, 102159 (2020).

44. Guo, K. et al. Transcription factor POU4F2 promotes colorectal cancer cell migration and invasion through hedgehogLmediated epithelialLmesenchymal transition. Cancer Science vol. 112 4176–4186 Preprint at 10.1111/cas.15089 (2021).

45. Feofanova, E. V. et al. Whole-Genome Sequencing Analysis of Human Metabolome in Multi-Ethnic Populations. Nat. Commun. 14, 1–12 (2023).

46. van der Flier, W. M. & Scheltens, P. Amsterdam Dementia Cohort: Performing Research to Optimize Care. J. Alzheimers. Dis. 62, 1091–1111 (2018).

47. Legdeur, N. et al. Resilience to cognitive impairment in the oldest-old: design of the EMIF-AD 90+ study. BMC Geriatr. 18, 289 (2018).

48. Boomsma, D. I. et al. Netherlands Twin Register: From Twins to Twin Families. Twin Res. Hum. Genet. 9, 849–857 (2006).

49. Konijnenberg, E. et al. The EMIF-AD PreclinAD study: study design and baseline cohort overview. Alzheimers. Res. Ther. 10, 75 (2018).

50. Albert, M. S. et al. The diagnosis of mild cognitive impairment due to Alzheimer’s disease: recommendations from the National Institute on Aging-Alzheimer’s Association workgroups on diagnostic guidelines for Alzheimer’s disease. Alzheimers. Dement. 7, (2011).

51. McKeith, I. G. et al. Diagnosis and management of dementia with Lewy bodies: Fourth consensus report of the DLB Consortium. Neurology 89, (2017).

52. McKhann, G. M. et al. The diagnosis of dementia due to Alzheimer’s disease: recommendations from the National Institute on Aging-Alzheimer’s Association workgroups on diagnostic guidelines for Alzheimer’s disease. Alzheimers. Dement. 7, (2011).

53. Rascovsky, K. et al. Sensitivity of revised diagnostic criteria for the behavioural variant of frontotemporal dementia. Brain 134, 2456 (2011).

54. Tijms, B. M. et al. Unbiased Approach to Counteract Upward Drift in Cerebrospinal Fluid Amyloid-β 1-42 Analysis Results. Clin. Chem. 64, 576–585 (2018).

55. Duits, F. H. et al. The cerebrospinal fluid ‘Alzheimer profile’: easily said, but what does it mean? Alzheimers. Dement. 10, 713–723.e2 (2014).

56. Tesi, N. et al. Centenarian controls increase variant effect sizes by an average twofold in an extreme case–extreme control analysis of Alzheimer’s disease. Eur. J. Hum. Genet. 27, 244–253 (2018).

57. Purcell, S. et al. PLINK: a tool set for whole-genome association and population-based linkage analyses. Am. J. Hum. Genet. 81, 559–575 (2007).

58. Quinlan, A. R. & Hall, I. M. BEDTools: a flexible suite of utilities for comparing genomic features. Bioinformatics vol. 26 841–842 Preprint at 10.1093/bioinformatics/btq033 (2010).

59. Giambartolomei, C. et al. Bayesian test for colocalisation between pairs of genetic association studies using summary statistics. PLoS Genet. 10, e1004383 (2014).

60. Benner, C. et al. FINEMAP: efficient variable selection using summary data from genome-wide association studies. Bioinformatics 32, 1493–1501 (2016).

61. Wang, G., Sarkar, A., Carbonetto, P. & Stephens, M. A simple new approach to variable selection in regression, with application to genetic fine mapping. J. R. Stat. Soc. Series B Stat. Methodol. 82, 1273–1300 (2020).

62. Weissbrod, O. et al. Functionally informed fine-mapping and polygenic localization of complex trait heritability. Nat. Genet. 52, 1355–1363 (2020).

