## Supplementary figures and images for "Quantitative trait loci mapping of circulating metabolites in cerebrospinal fluid to uncover biological mechanisms involved in brain-related phenotypes"

### Fig_S1_mQTL_replication_biofluids_2023-03-14.jpg

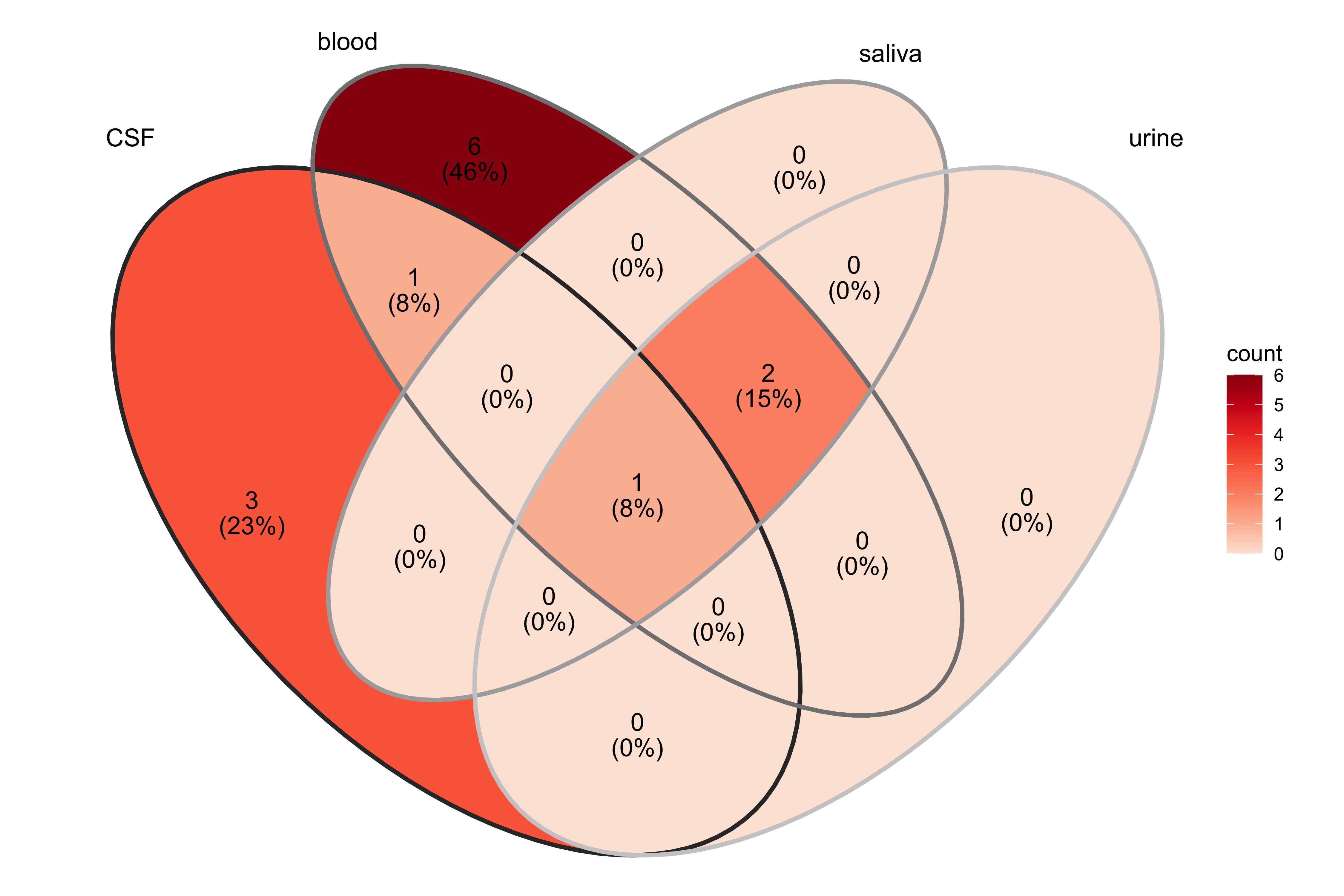

### Fig_S2_N.epsilon.-Dimethyl-L-lysine_rs10883083_33_2023-02-14.jpg

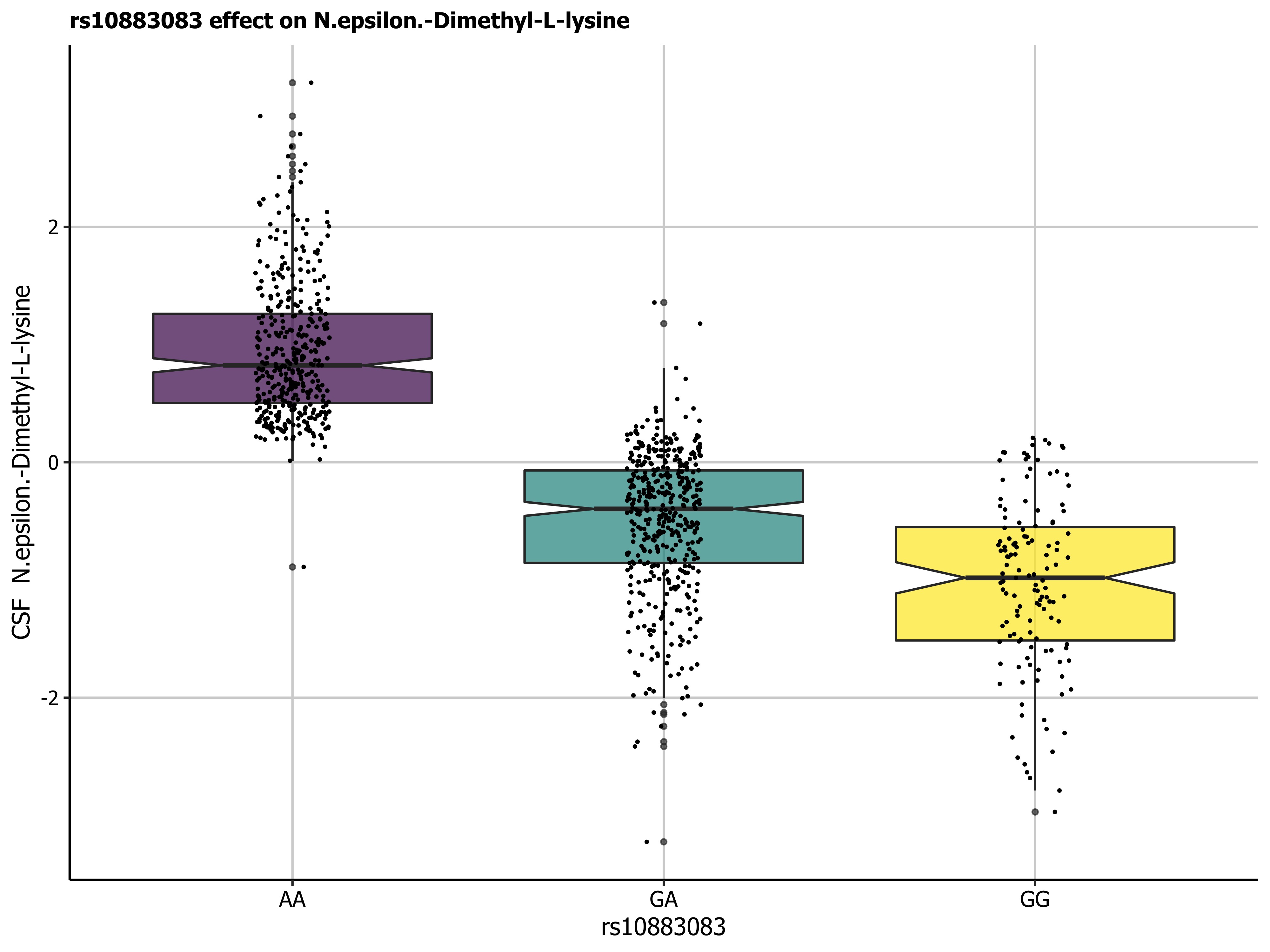

### Fig_S3_7.25_144.10_chr10_98392298_C_G_70_2023-02-14.jpg

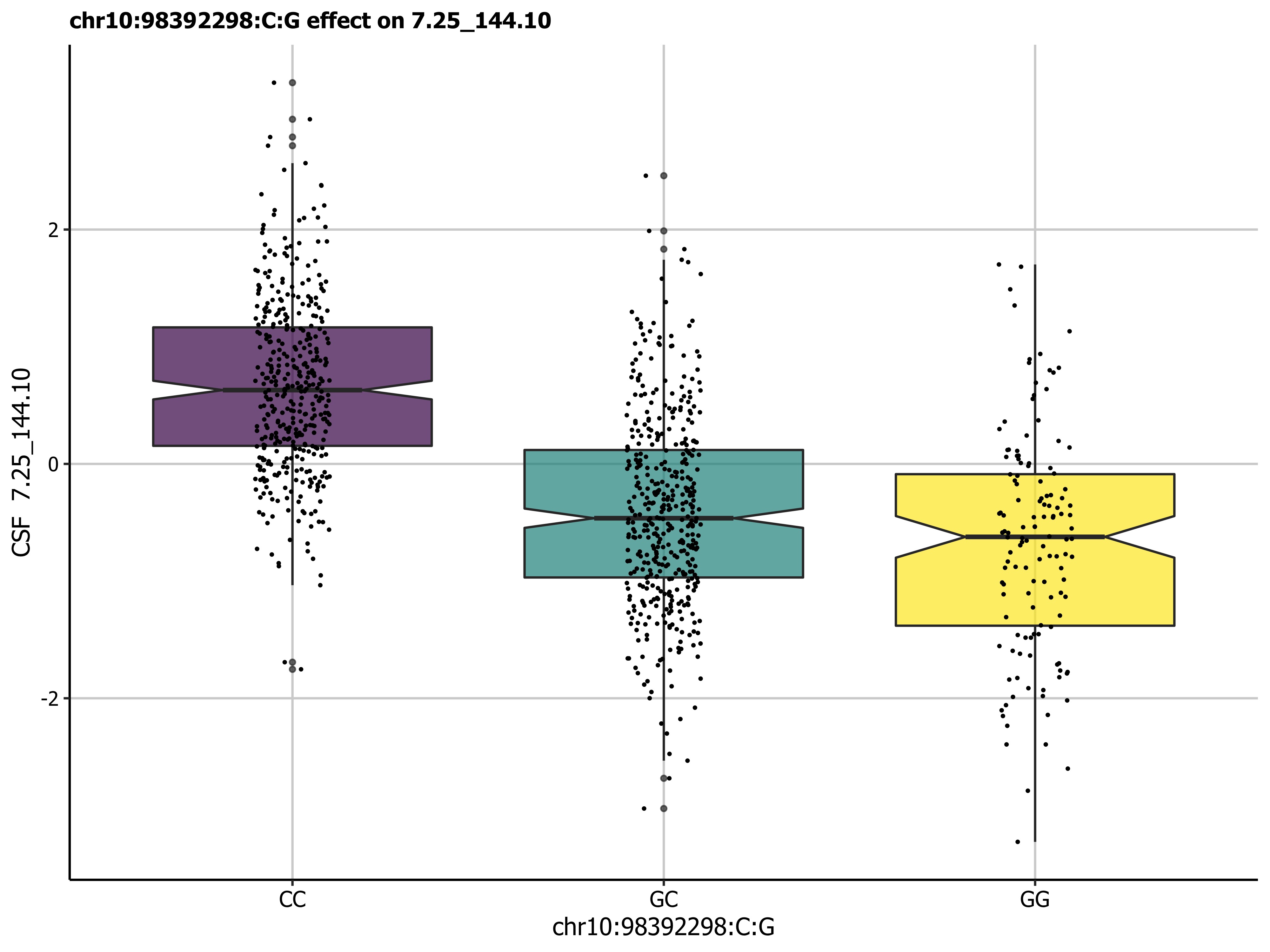

### Fig_S4_9.29_260.20_chr10_98392298_C_G_58_2023-02-14.jpg

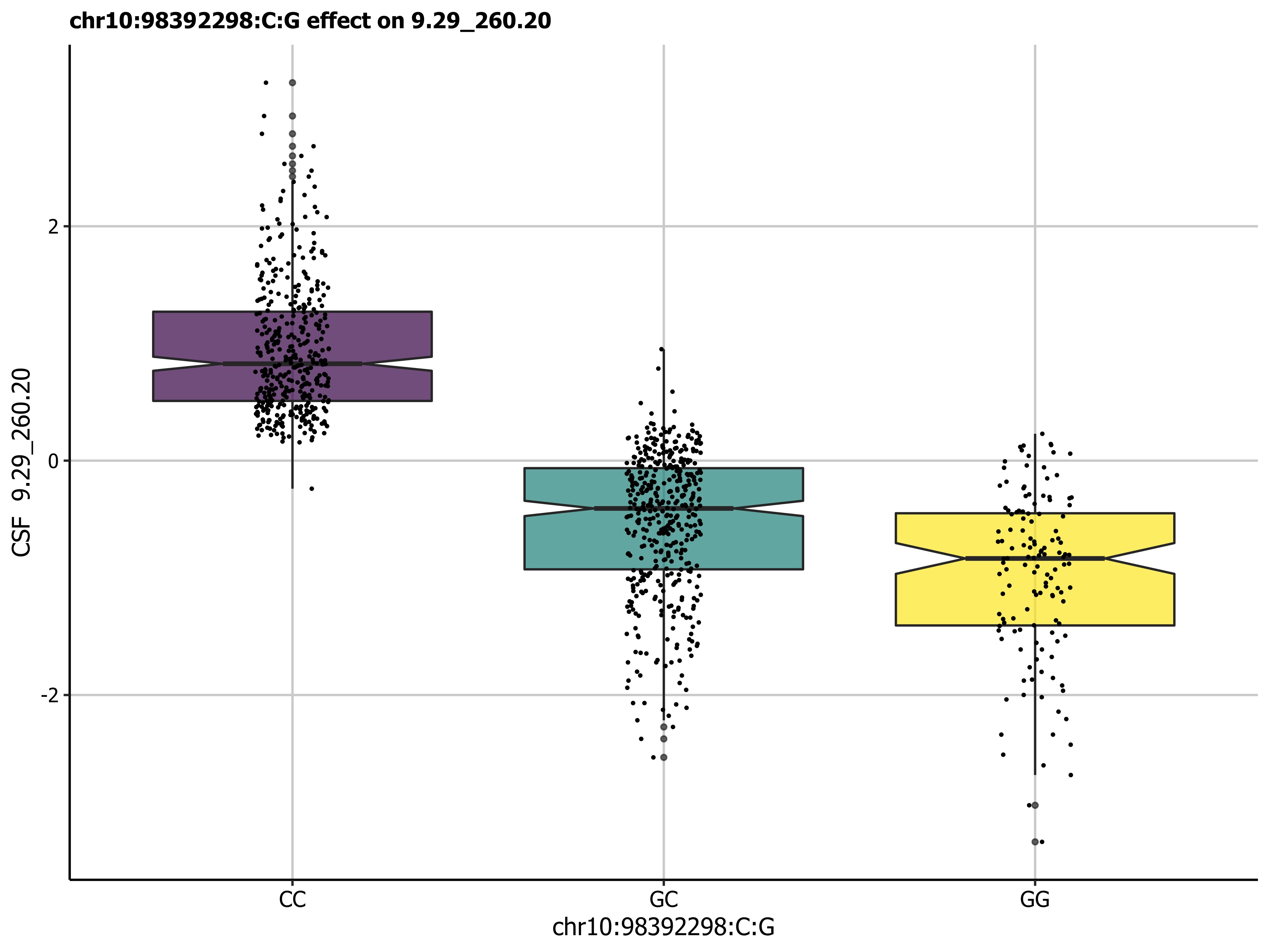

### Fig_S5_9.30_130.09_rs10883083_116_2023-02-14.jpg

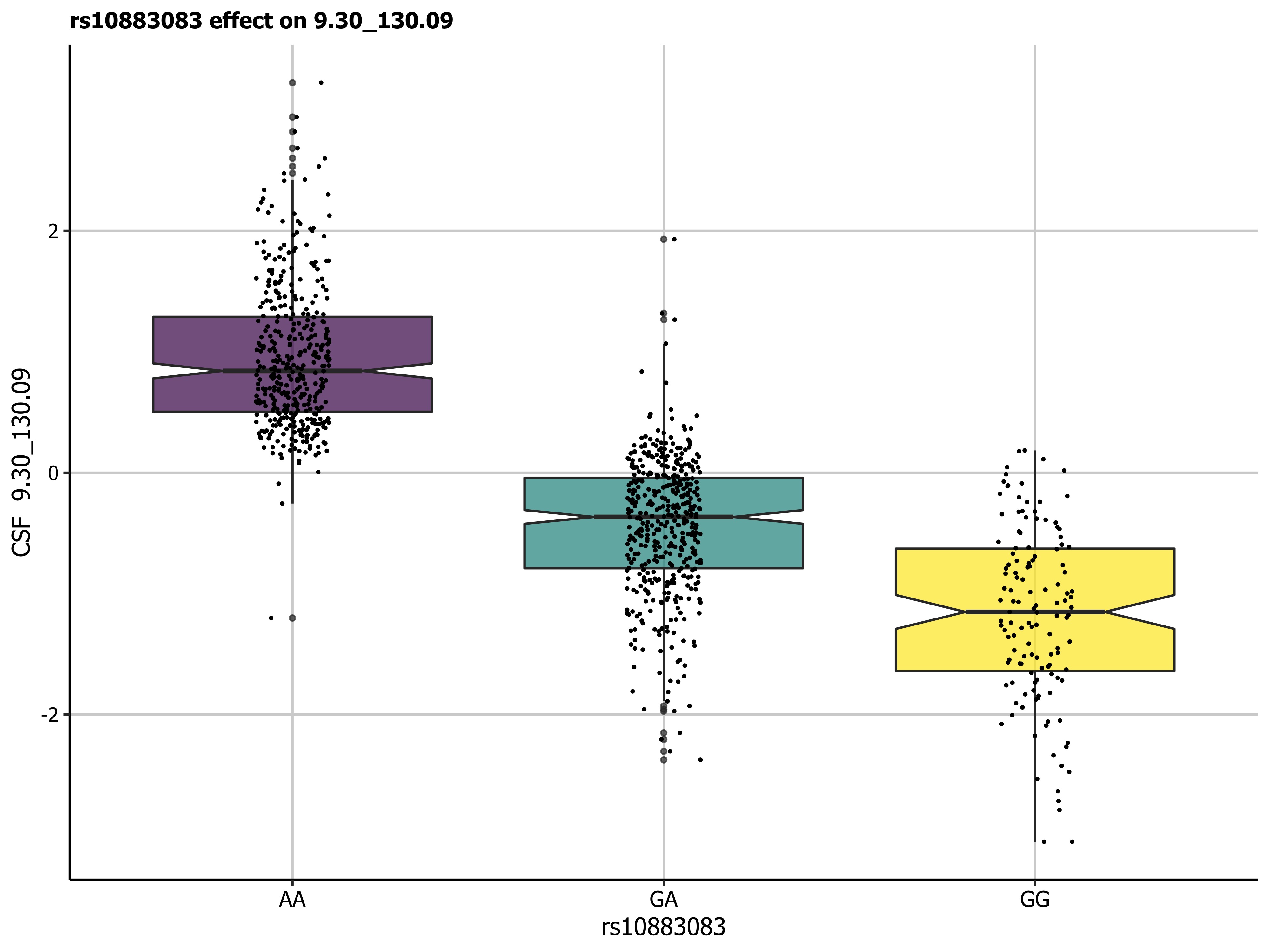

### Fig_S6_9.31_161.13_rs10883083_100_2023-02-14.jpg

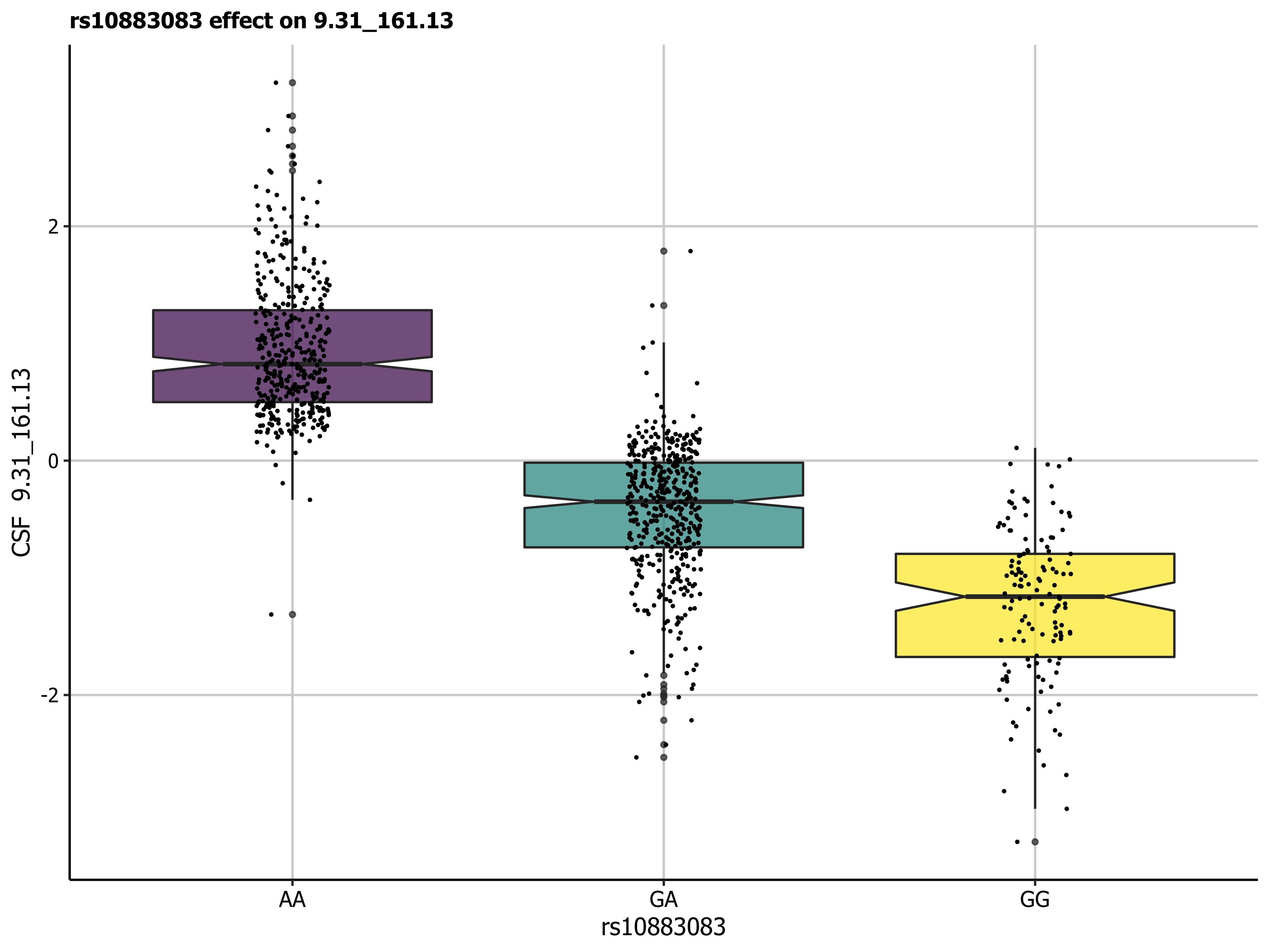

### Fig_S7_N-Acetylhistidine_rs740104_25_2023-02-14.jpg

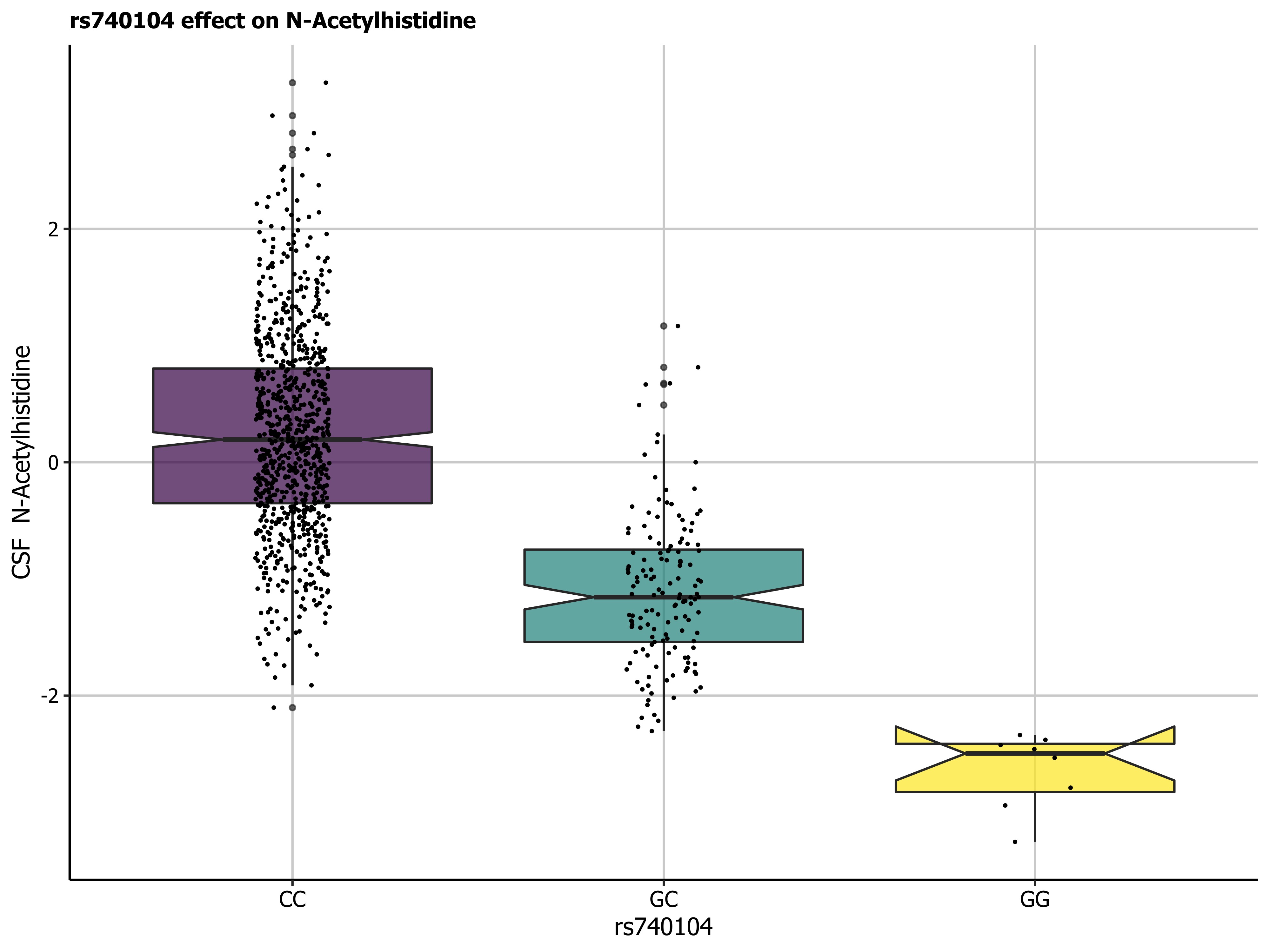

### Fig_S8_8.10_220.07_rs740104_84_2023-02-14.jpg

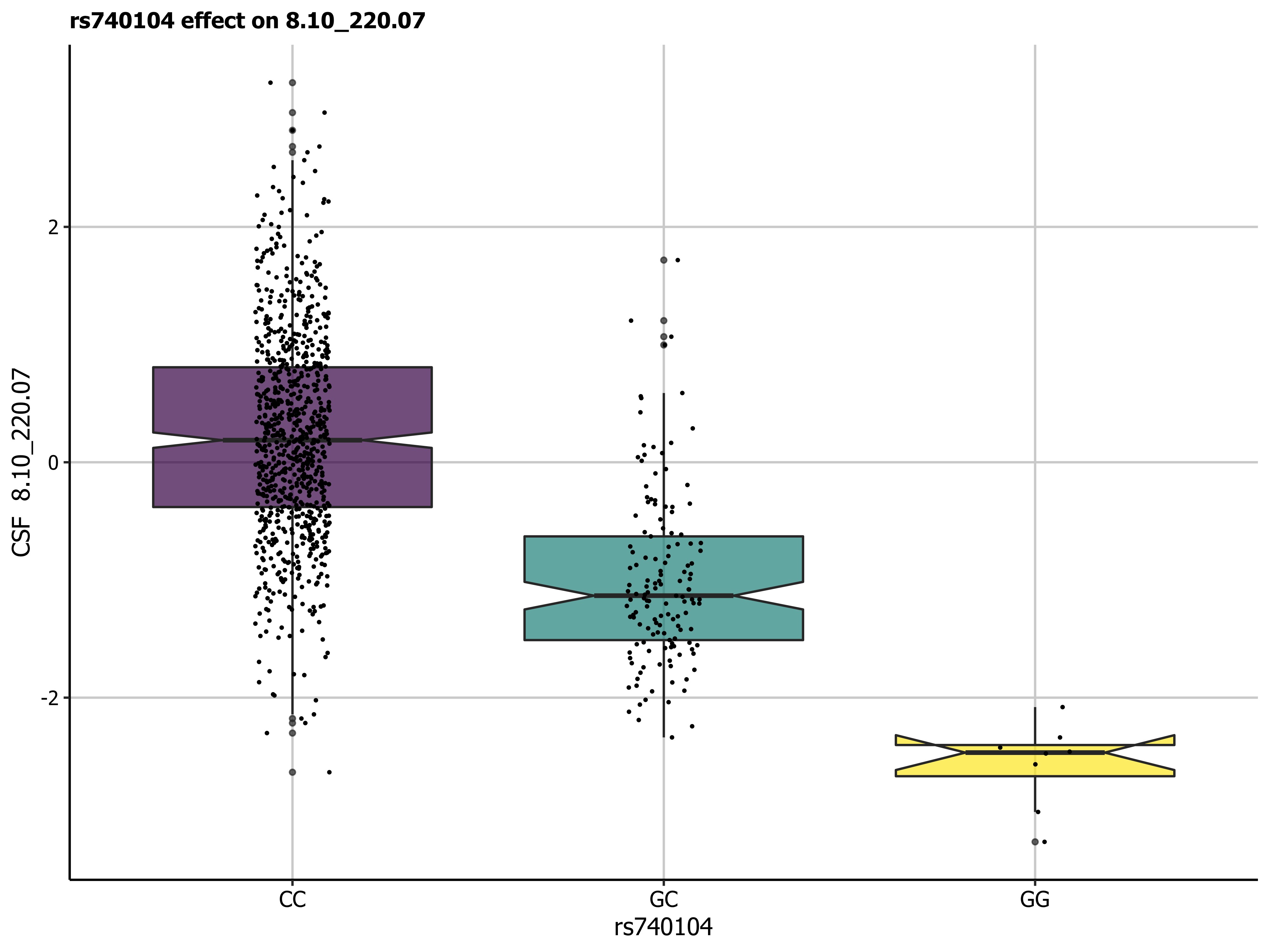

### Fig_S9_7.40_268.13_rs740104_80_2023-02-14.jpg

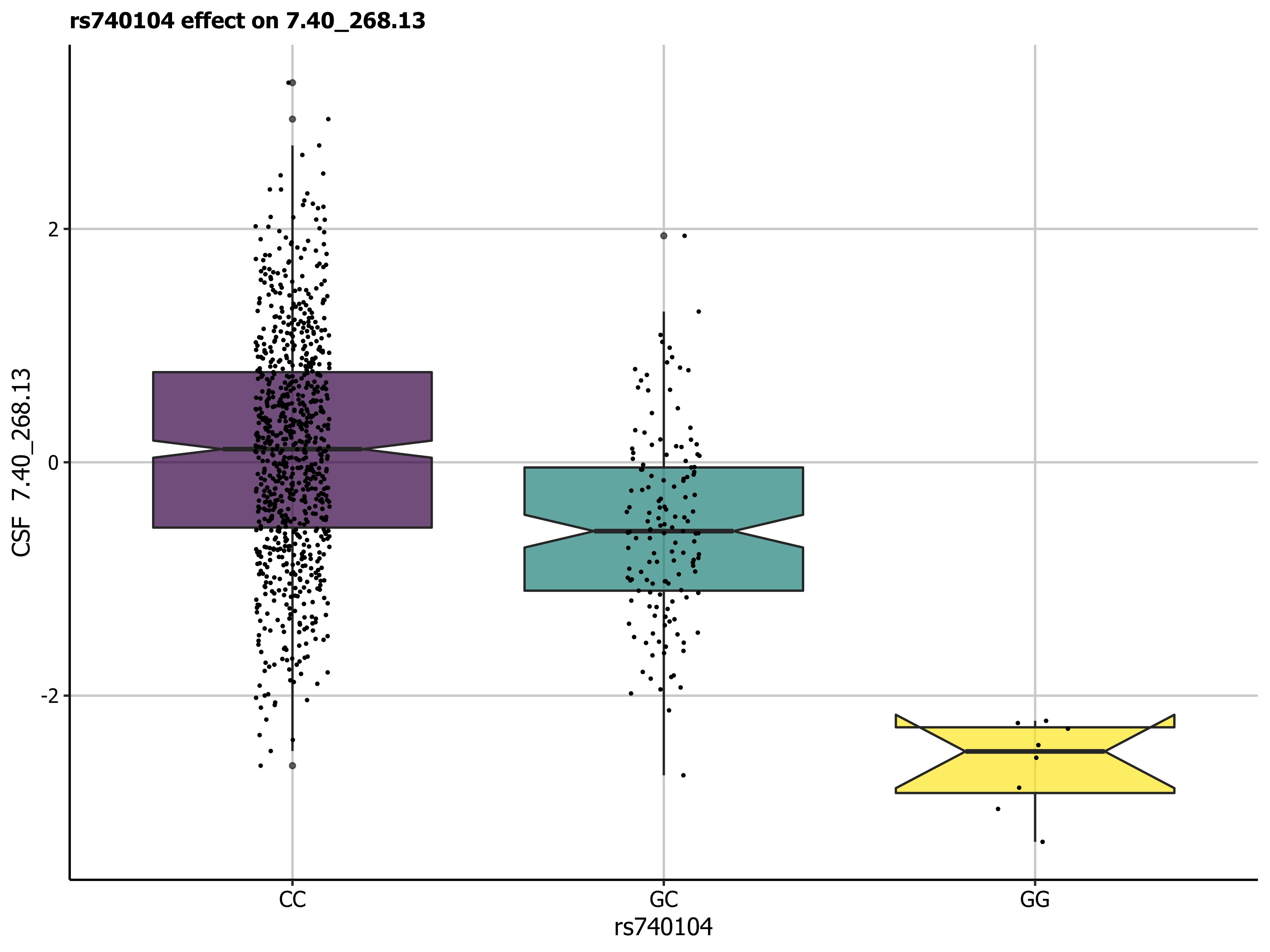

### Fig_S10_Manhattan_Hypaphorine.tiff

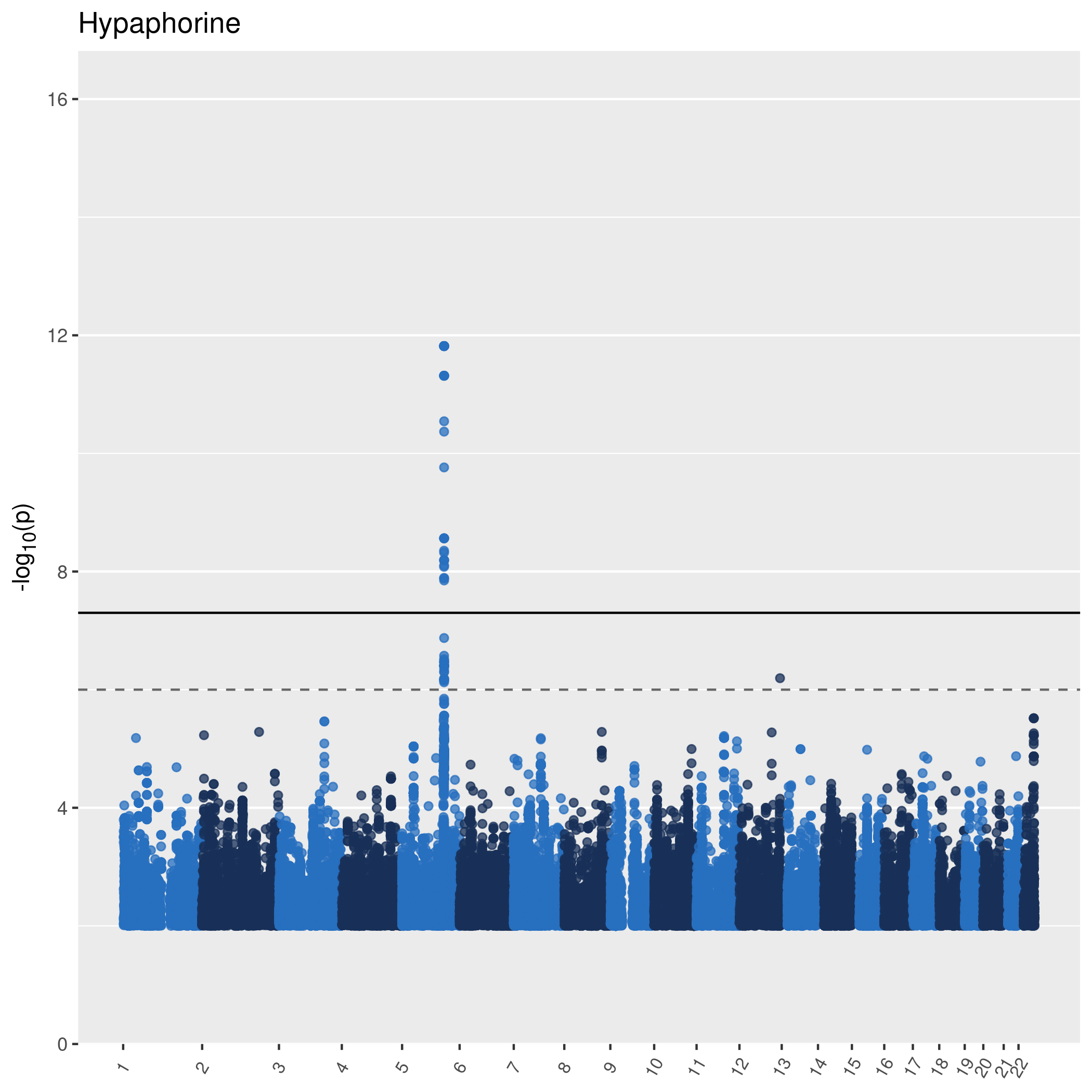

### Fig_S11_Hypaphorine_chr5_132389258_T_C_31_2023-02-14.jpg

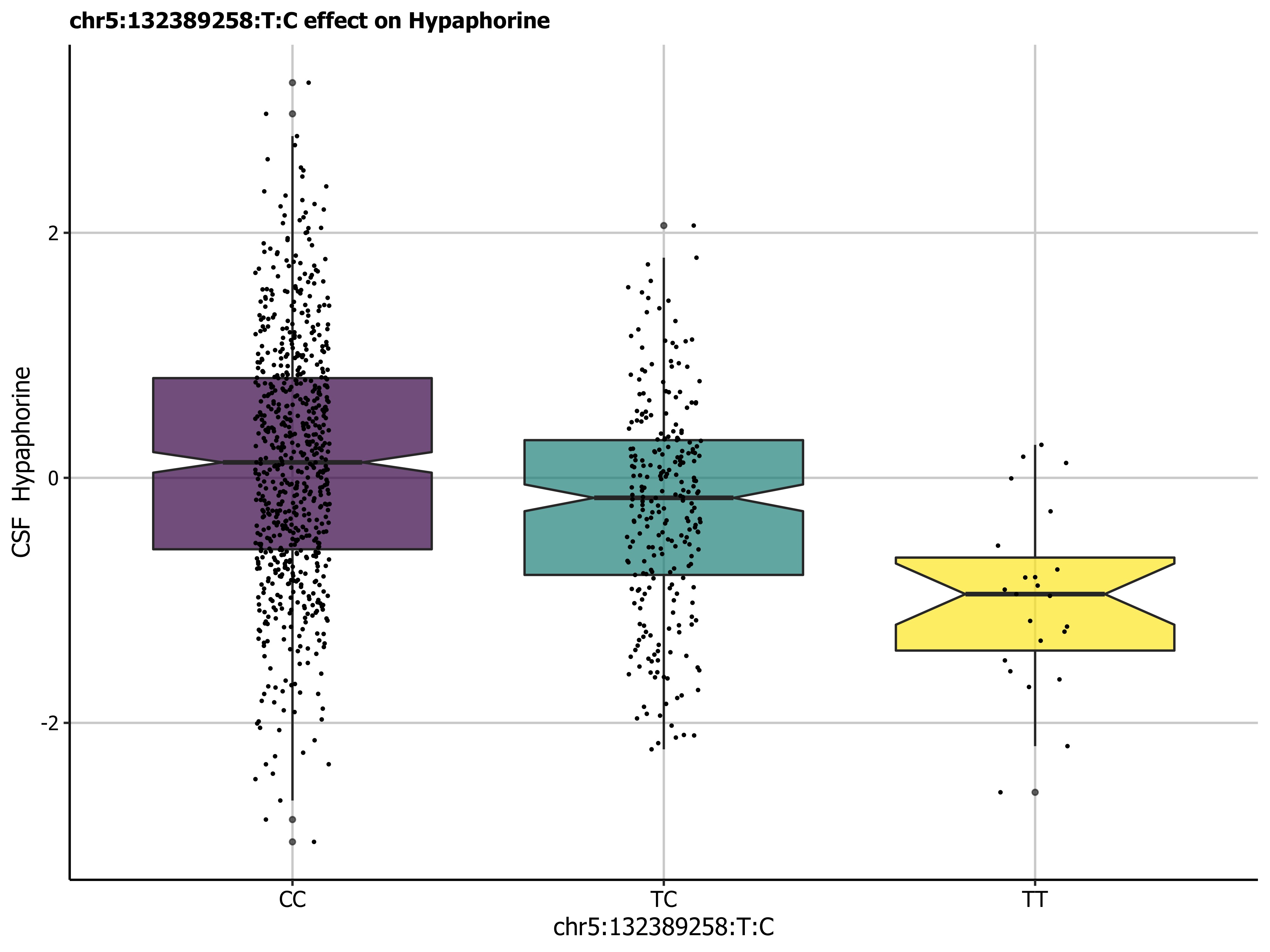

### Fig_S12_MWAS_weights_platform_2023-01-24.tiff

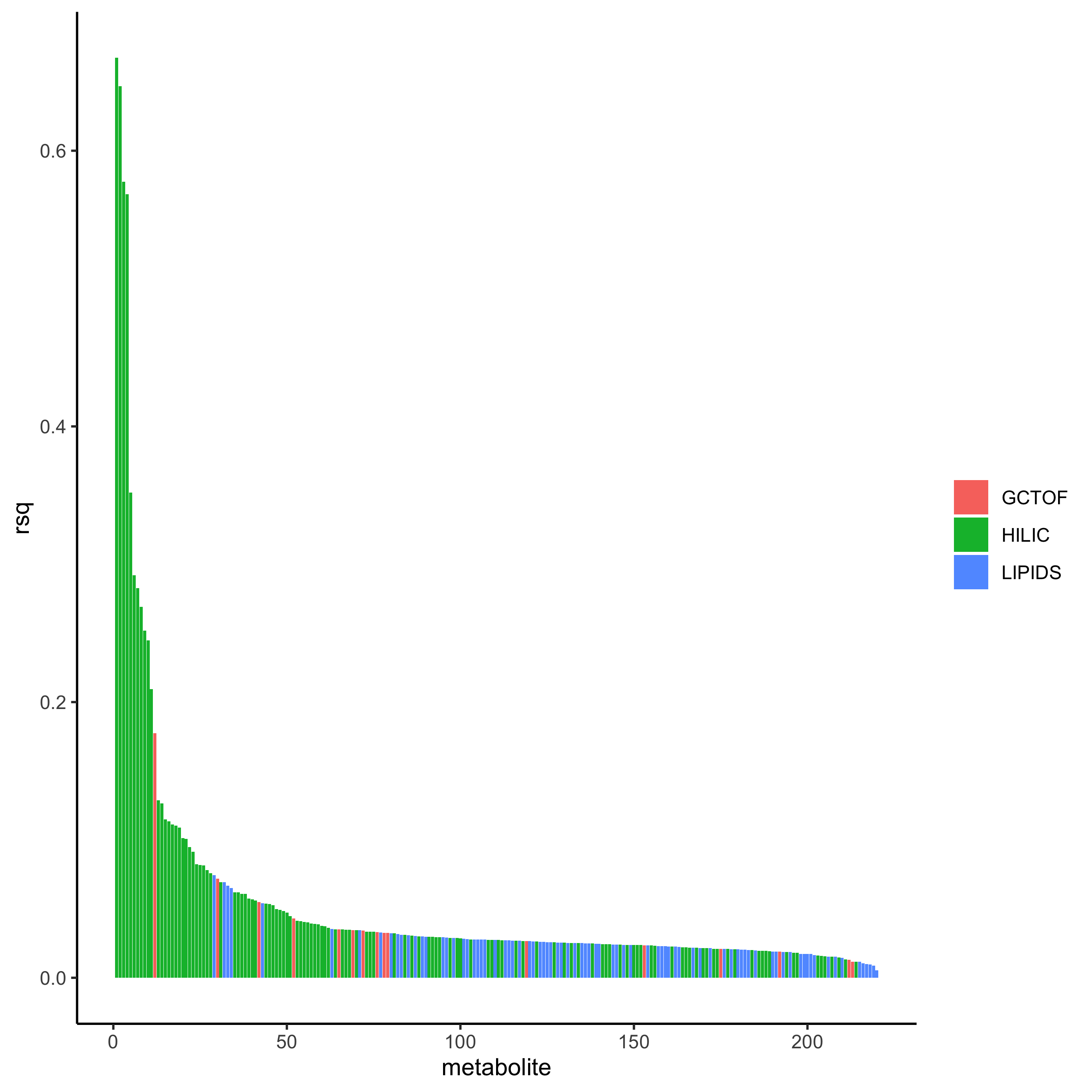

### Fig_S13_MWAS_weights_2023-01-24.tiff

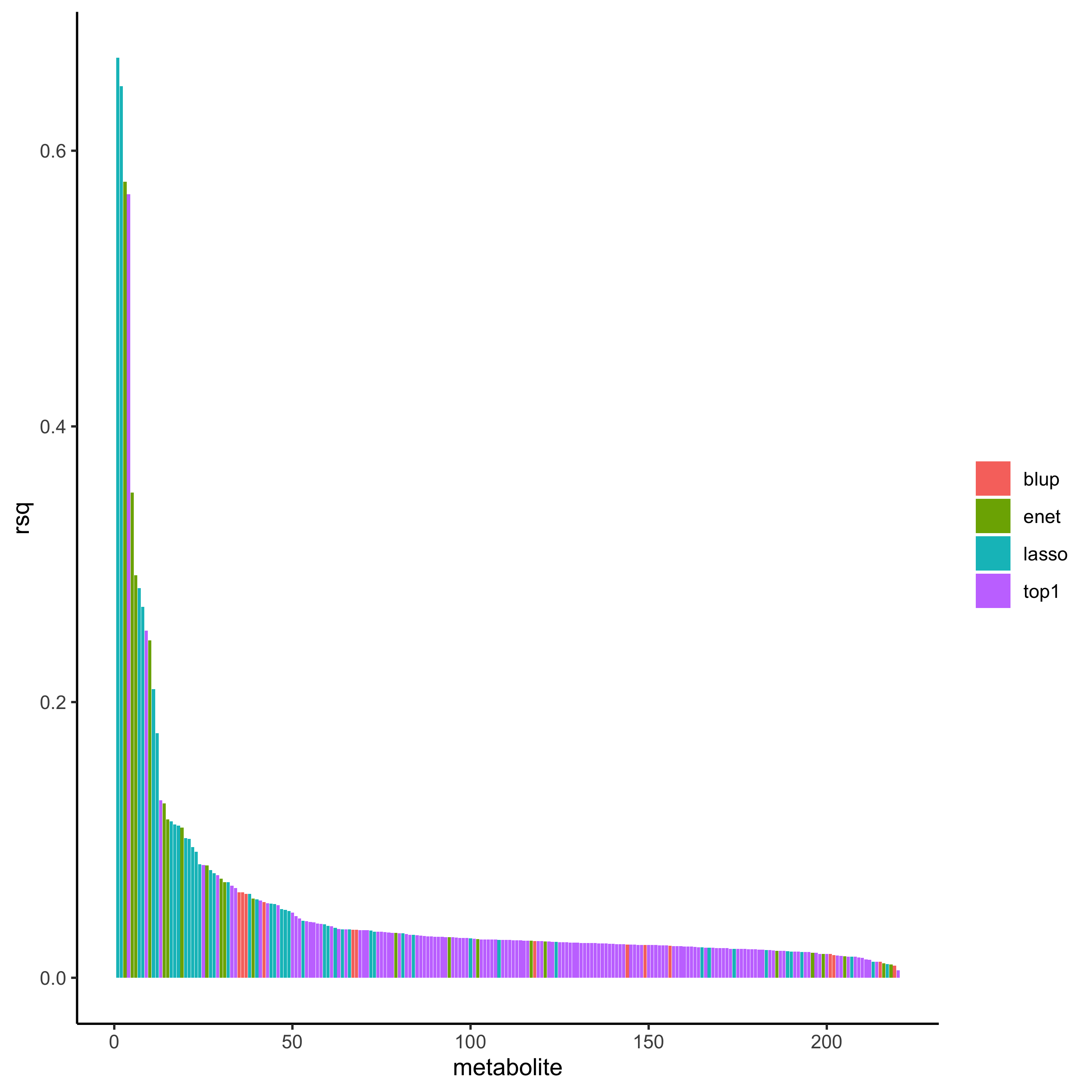

### Fig_S14_TWAS_Manhattan_2023-06-19.tiff

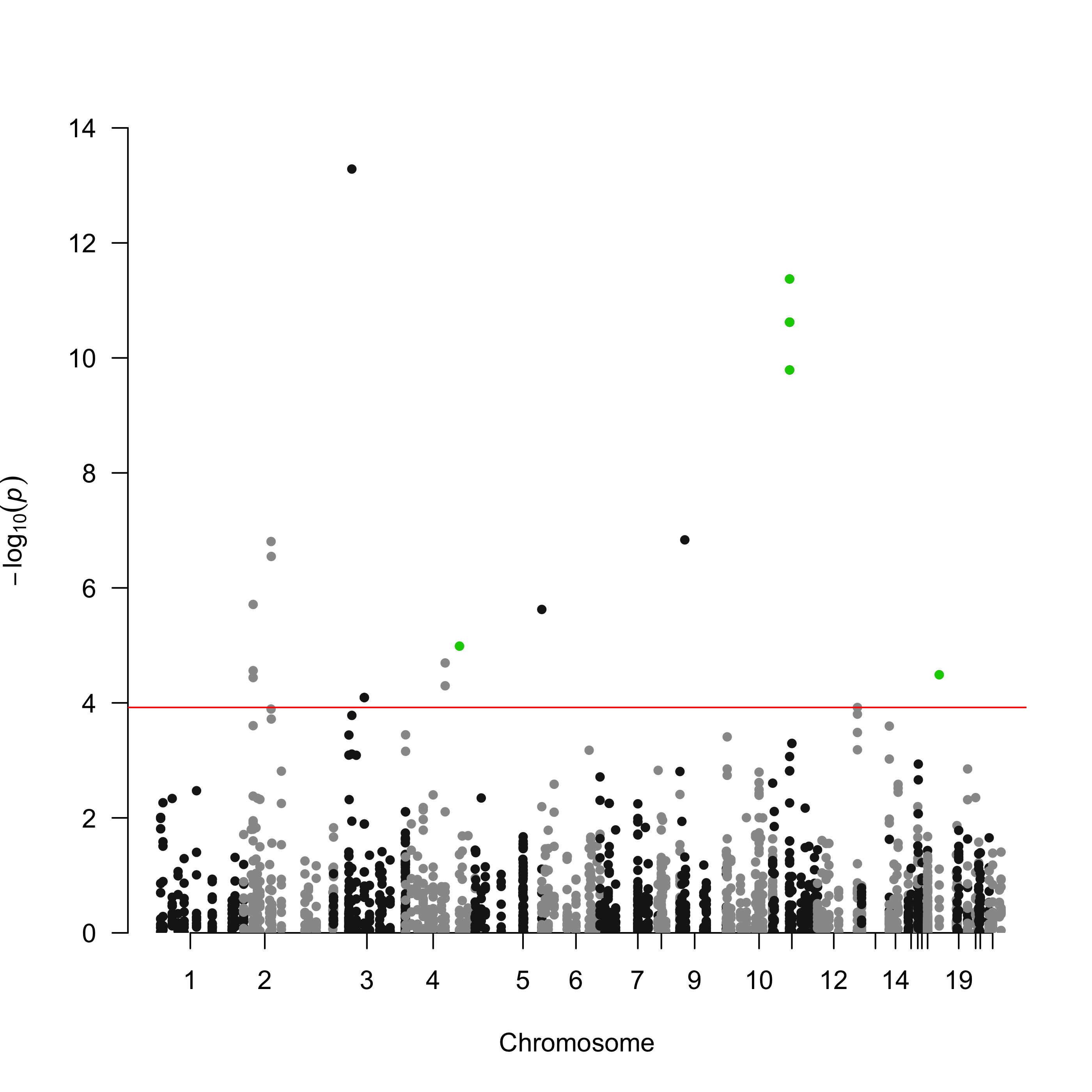

### Fig_S15_Bicalutamide_rs1155168_22_2023-02-14.jpg

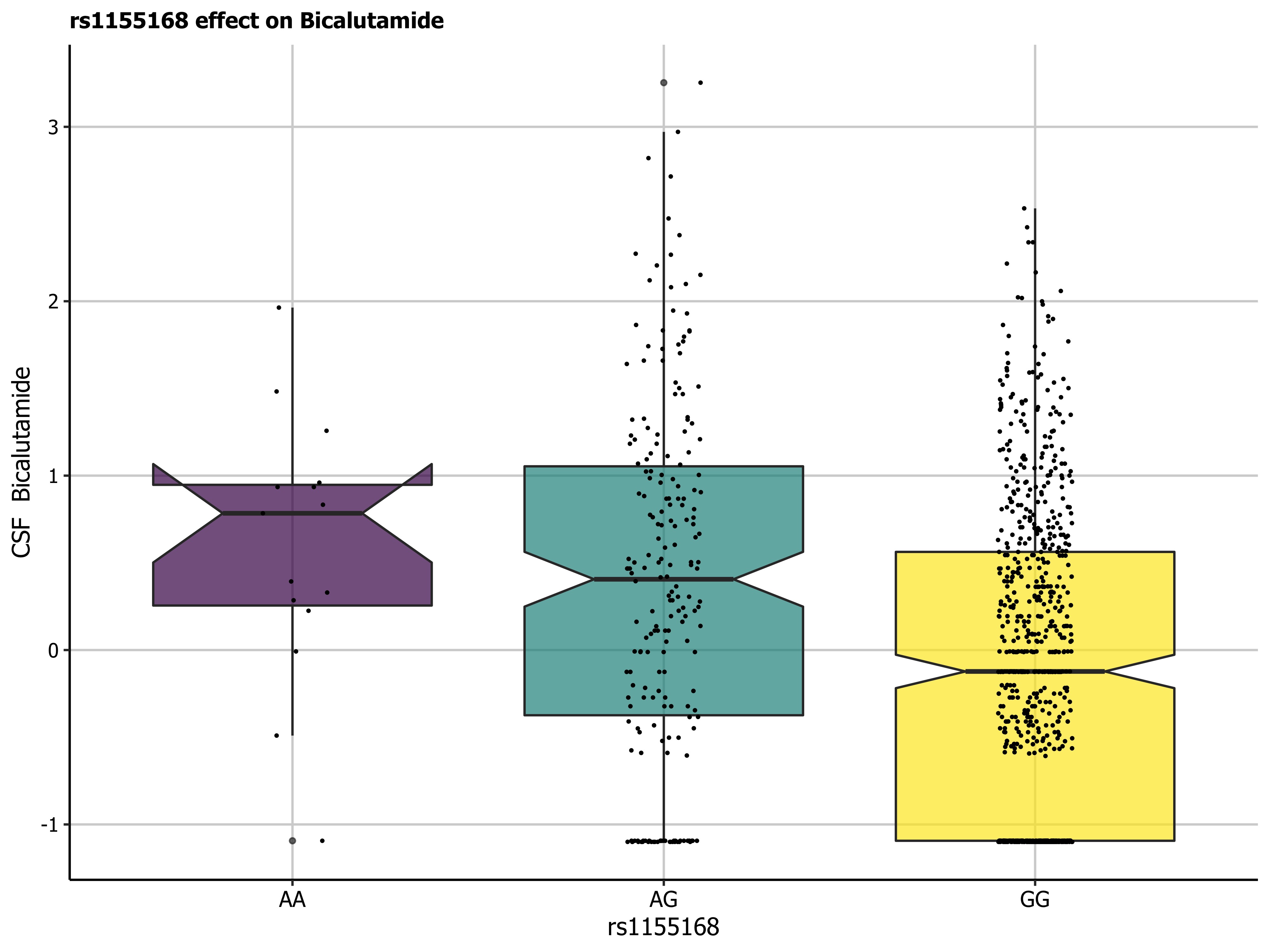
